# Large oncosomes reprogram bone marrow mesenchymal stem cells to establish a pre-metastatic niche

**DOI:** 10.64898/2026.08.20.746089

**Authors:** Taylon Felipe Silva, Virginia Marcia Concato-Lopes, Fatima Bedier, Lauren Newman, Diana Kitka, Jamelle Brown, Blandine Victor, Minhyung Kim, Tatyana Vagner, Catherine Grasso, Dmitriy Sheyn, Sungyong You, Michael R. Freeman, Fayyaz S. Sutterwala, Helen S. Goodridge, Caroline Jefferies, Paola de Candia, Jlenia Guarnerio, Dolores Di Vizio

## Abstract

Metastatic progression depends on the systemic remodeling of distant tissues before tumor cell arrival, yet the cancer-derived signals that orchestrate this process remain poorly understood. Large oncosomes (LOs) are atypically large (>1 µm), tumor-derived extracellular vesicles shed by invasive cancer cells. Here, we demonstrate that LOs function as systemic mediators of pre-metastatic niche formation by activating innate immune sensing in bone marrow mesenchymal stem cells (BM-MSCs). Systemic administration of prostate- and breast cancer-derived LOs to immunocompetent tumor-bearing mice did not affect primary tumor growth but increased metastatic burden. LOs induced a robust, dose-dependent interferon-driven inflammatory program, characterized by interferon-stimulated genes and neutrophil chemokines. Mechanistically, this response was driven by LO-associated nucleic acids activating convergent cytosolic DNA- and RNA-sensing pathways in recipient stromal cells. LO-conditioned BM-MSCs promoted the accumulation and polarization of neutrophils toward an immunosuppressive, polymorphonuclear myeloid-derived suppressor cell (PMN-MDSC) phenotype. *In vivo*, genetic suppression of LO shedding in tumor cells reduced PMN-MDSC accumulation in the bone marrow, which was reverted by systemic LO administration. Together, these findings establish LOs as specialized carriers of immunomodulatory signals that reprogram the bone marrow microenvironment to support metastatic colonization, identifying a previously unrecognized mechanism linking tumor vesiculation to immune remodeling at distant sites.

**Graphical Abstract:** Large oncosomes systemically reprogram bone marrow mesenchymal stem cells through innate nucleic acid sensing, driving inflammatory and immunosuppressive remodeling that supports pre-metastatic niche formation, revealing a novel tumor-host communication axis in cancer.

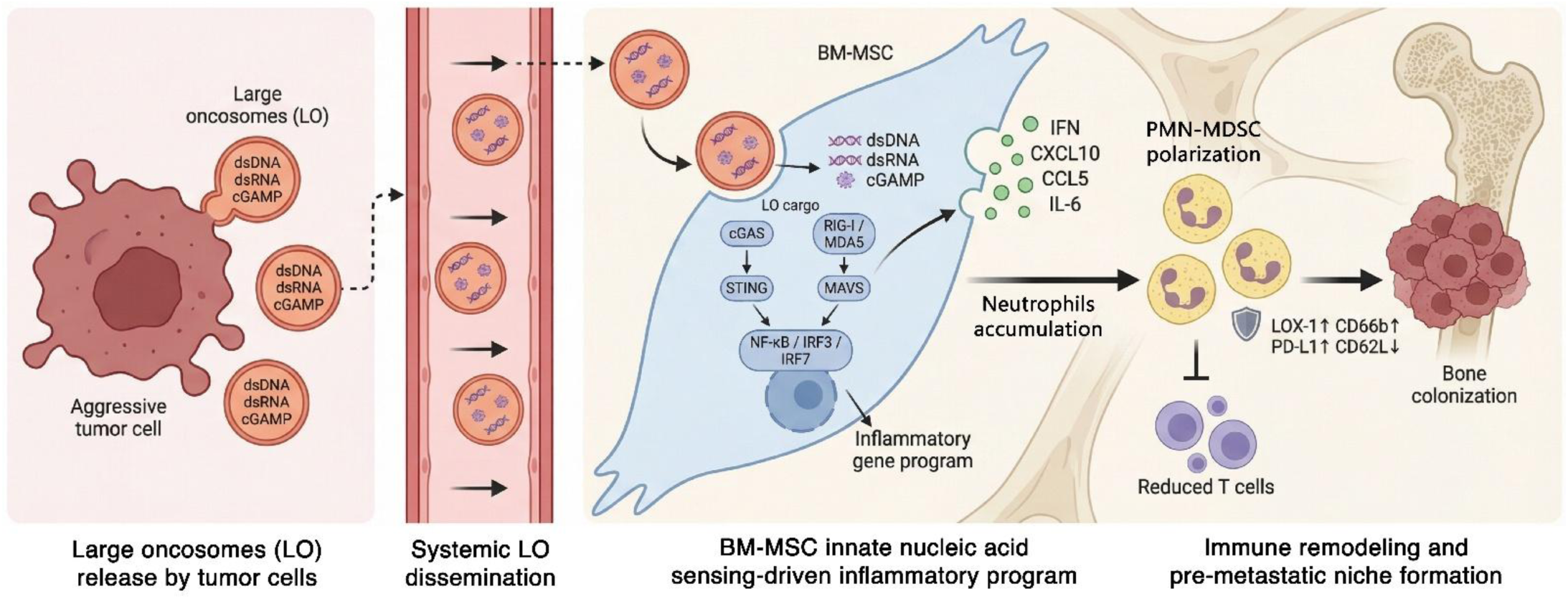

## INTRODUCTION

Metastatic progression is increasingly recognized as a systemic process driven by continuous communication between tumor cells and distant host tissues^[1–3]^. Through this reciprocal crosstalk, primary tumors actively remodel metastatic sites even before the arrival of disseminated tumor cells, creating permissive microenvironments that support metastatic colonization^[3–5]^. Extracellular vesicles (EVs) have emerged as central mediators of this long-range communication, coordinating diverse processes that include stromal activation, immune modulation, and organ-specific niche formation^[4,6,7]^. The expanding recognition of EV heterogeneity has further highlighted the need to define how distinct EV populations contribute to these processes^[8–10]^.

Large oncosomes (LOs) represent a biologically distinct class of tumor-derived EVs. Unlike exosomes and other small EVs (sEVs), LOs are atypically large (>1 μm), membrane bleb-derived vesicles selectively shed by highly migratory and invasive tumor cells with metastatic capacity^[11–13]^. Their biogenesis is tightly linked to cytoskeletal remodeling, non-apoptotic, amoeboid membrane blebbing, and cellular programs associated with nuclear shape instability and highly malignant tumor phenotypes^[11,14,15]^. LOs are abundant in the circulation of patients with advanced malignancies^[9,11,16–19]^ and have been implicated in multiple aspects of tumor progression, including tumor cell adhesion, invasion, angiogenesis and metastatic dissemination^[11,20–23]^.

The biological function of LOs is closely linked to their unique molecular composition. Profiling studies including comparative multiomics of LOs and S-EVs have revealed that LOs are enriched in diverse classes of tumor-derived cargo, such as proteins, lipids, genomic DNA, and multiple RNA species^[9,12,16,21,24–27]^. The abundance and diversity of nucleic acids within LOs suggest that these vesicles may serve as more efficient vehicles for the systemic dissemination of tumor-derived genetic material in comparison with sEVs. A recent study have further implicated LOs in pre-metastatic niche conditioning and organ-specific metastatic tropism, extending their role beyond local tumor progression to systemic regulation of metastatic competence^[22]^.

Tumor-derived nucleic acids have emerged as potent regulators of stromal and immune cell behavior. In both cancer and tissue injury, aberrant DNA and RNA species activate innate immune sensing pathways that converge on type I interferon as an inflammatory signaling program. This type of response can profoundly influence stromal cell states, myeloid cell recruitment, and the establishment of immunoregulatory microenvironments that support tumor progression and metastasis ref. Cancer-derived molecular signals, including mitochondrial DNA and other damage-associated nucleic acids, have been shown to reprogram myeloid populations through innate immune mechanisms^[28]^. Similarly, LOs isolated from cancer stem cells have been reported to carry cytokines and other bioactive cargo capable of modulating myeloid cell behavior^[29]^. Together, these observations raise a fundamental question on the biology of metastasis: how are tumor-derived inflammatory signals disseminated systemically to coordinate stromal and immune remodeling across distant tissues?

In this study we investigated how tumor-derived LOs contribute to systemic communication during metastatic progression using prostate cancer and breast cancer as two models of bone metastasis. Using integrated molecular, cellular, and *in vivo* approaches, we defined the mechanisms by which LO-associated nucleic acids activate specific innate immune pathways in bone marrow stromal cells, thereby promoting stromal and immune reprogramming linked to metastasis. Together, this work provides a novel mechanistic framework for understanding how LOs mediate tumor-host communication during metastasis.

## MATERIALS AND METHODS

### Cell culture and primary cell preparation

The human prostate cancer cell line PC3 and human breast cancer cell line MDA-MB-231 were maintained in Dulbecco’s Modified Eagle Medium (DMEM; GIBCO). The murine breast cancer cell line 4T1.2 was maintained in alpha Minimum Essential Medium (alpha-MEM; GIBCO). Human primary bone marrow-derived mesenchymal stem cells (BM-MSCs) (ATCC) were cultured in MesenPRO RS medium (GIBCO) according to the manufacturer’s recommendations. Murine bone marrow cells and BALB/c-derived OriCell mouse BM-MSCs (Cyagen) were maintained in OriCell Complete Medium for Mouse BM-MSCs (Cyagen). Unless otherwise indicated, media were supplemented with 10% fetal bovine serum, 2 mM L-glutamine, and 1% penicillin-streptomycin. Cells were cultured at 37°C in a humidified incubator with 5% CO2. Primary murine bone marrow cells were isolated from long bones of WT B6;129SF2/J mice and genetically modified strains including B6;129-Mavs^tm1Zjc^/J and Sting1^gt^ mice. Femurs and tibiae were aseptically harvested, cleaned of adherent tissue, and flushed with pre-warmed culture medium using a sterile syringe. Cell suspensions were filtered through a 70 µm cell strainer, red blood cells were lysed using ammonium-chloride-potassium (ACK) lysis buffer, and cells were washed and resuspended in the appropriate culture medium. Primary human neutrophils were isolated from freshly collected peripheral blood from healthy donors using the EasySep Direct Human Neutrophil Isolation Kit according to the manufacturer’s protocol and maintained in RPMI-1640 medium. Human primary neutrophils-related studies were conducted under Cedars-Sinai Institutional Review Board approval IRB 19627 and in accordance with institutional and ethical guidelines. All human and murine cell lines were obtained from ATCC unless otherwise indicated, and primary BM-MSCs were obtained commercially and used at low passage. Cell lines were used at a maximum passage of 12, and primary cells were used at passage ≤6. Cultures were routinely monitored for mycoplasma contamination using MycoStrip Mycoplasma Detection Kit (InvivoGen) and/or MycoAlert PLUS (Lonza). Cultures used for EV production had at least 98% viability before conditioned medium collection, as assessed by trypan blue exclusion or equivalent viability testing.

### Mouse models

All animal experiments were approved by the Cedars-Sinai Institutional Animal Care and Use Committee under protocol IACUC008508 (PI: Di Vizio) and were performed in accordance with institutional guidelines for animal care and use. Mice were purchased from The Jackson Laboratory and used between 6 and 8 weeks of age. Male mice were used for prostate cancer and female mice for breast cancer, as appropriate for each experimental model. BALB/cJ mice were used for syngeneic orthotopic 4T1.2 breast cancer models, intratibial BM-MSC priming experiments, and LO rescue studies. C57BL/6J mice were used for bone marrow stromal and immune recruitment assays when indicated. WT B6;129SF2/J mice, Sting1^gt^ mice, and B6;129-Mavs^tm1Zjc^/J mice were used for genetic validation of cytosolic nucleic acid-sensing pathways in primary bone marrow cultures. Animals were housed under standard pathogen-free conditions with *ad libitum* access to food and water and were monitored routinely according to approved veterinary and IACUC guidelines.

### *In vivo* BM-MSC priming and intracardiac breast cancer metastasis model

BALB/c-derived mouse BM-MSCs (Cyagen OriCell) were pretreated with nLuc-GFP 4T1.2-derived LOs, sEVs, or vehicle control and injected into the tibia of syngeneic female BALB/c mice. A total of 1×10^5^ BM-MSCs in 10 uL were injected per tibia. Forty-eight hours later, 5×10^4^ GFP/NanoLuc reporter 4T1.2 cells in 100 uL PBS were injected into the left cardiac ventricle. Mice were monitored for 2 weeks by IVIS imaging using Nano-Glo Fluorofurimazine *In vivo* Substrate Kit (Promega) to quantify metastatic spread and focal tibial lesions. All mice were female and 6-8 weeks old.

### Orthotopic 4T1.2 tumor education and breast cancer metastasis model

For systemic EV education during primary tumor progression, female 6-8-week-old BALB/c mice were orthotopically implanted with 2×10^5^ nLuc-GFP 4T1.2 cells into the lower right mammary fat pad in Corning Matrigel growth factor reduced basement membrane matrix, phenol red-free. Mice were assigned to vehicle, sEV, or LO education groups (15 animals per group). During the 14-day primary tumor growth phase, mice received four retro-orbital EV injections split across the 2-week interval, each consisting of 10 ug EV protein in 100 uL PBS. Primary tumors were surgically excised on day 14. Metastatic progression was monitored for an additional 60 days, and mice were euthanized at approximately day 75 or earlier if humane endpoints were reached. Primary tumor growth was measured during the initial phase, and metastatic burden was tracked by IVIS using Nano-Glo Fluorofurimazine *In vivo* Substrate Kit (Promega) and lesion analysis.

### Hollow fiber bioreactor culture

For large-scale EV production, selected cancer cell lines (PC3 and 4T1.2) were expanded in a hollow fiber bioreactor using C2018 cartridges (FiberCell Systems) with a 20 kDa molecular weight cutoff. Cells were seeded into the extracapillary space without prior coating at an initial target input of approximately 1 x 10^9^ cells. Cultures were maintained under continuous perfusion in high-glucose DMEM, and fetal bovine serum was gradually replaced with CDM-HD according to the manufacturer’s recommendations, until reach serum free condition and 10% CDM-HD. Bioreactor cultures were monitored by glucose consumption and maintained under high-density conditions, with glucose uptake targeted at approximately 1-1.5 g/day. Extracapillary space medium was collected every other day for EV enrichment. Bioreactors were maintained for up to 60 days, with culture health monitored by glucose consumption, absence of overt cell contamination/dead cells in conditioned medium, and confirmation of EV markers after isolation. Cultures were routinely monitored for mycoplasma.

### Enrichment and purification of LOs and sEVs from conditioned medium

EV enrichment was performed from conditioned medium collected from conventional 2D cultures or hollow fiber bioreactor cultures, following an adapted differential centrifugation and iodixanol density-gradient purification workflow. For 2D culture, cells were plated in 150 mm tissue-culture dishes (typically n=18 dishes per preparation) and expanded in serum-containing medium for at least 48 h. When cultures reached approximately 80-90% confluence, cells were washed twice with PBS and incubated in serum-free medium for 24 h before collection. Conditioned medium was cleared by three sequential centrifugation steps at 300 x g for 5 min each at 4 °C to remove cells and large debris. The LO-enriched fraction was then generated from the medium-speed pellets by centrifuging the cleared conditioned medium at 10,000 x g. The supernatant was subsequently centrifuged at 100,000 x g for 60 min to collect the sEV-enriched fraction. All ultracentrifugation steps were performed at 4°C using an SW28 swinging-bucket rotor (Beckman-Coulter, USA) where applicable. Fresh EV pellets were further resuspended in 0.2 µm-filtered PBS and loaded at the bottom of Beckman Coulter ultracentrifuge tubes (Cat. #344058). Discontinuous iodixanol gradients were prepared by sequentially layering 30% (4.3 mL; 1.20 g/mL), 25% (3 mL; 1.15 g/mL), 15% (2.5 mL; 1.10 g/mL), and 5% (6 mL; 1.08 g/mL) iodixanol solutions. Gradients were centrifuged at 100,000 x g for 4 h, at 4°C. LO fractions were recovered from the 1.10-1.15 g/mL density range, whereas sEVs derived from the 100,000 x g pellet were recovered from the 1.10 g/mL density fraction following previously standardized procedures^[9]^. Purified EVs were washed in 0.2 um-filtered PBS by ultracentrifugation at 100,000 x g for 60 min, resuspended in 0.2 um-filtered PBS, aliquoted and stored at −80 C. EV preparation aliquots were used only once after the first thaw and discarded afterwards, to avoid repeated freeze-thaw cycles. Unless otherwise stated, EV input for *in vitro* treatments was normalized by protein concentration at 20 µg/mL. Dose-response experiments used the EV concentrations indicated in the relevant figure legends. For *in vivo* EV education or rescue experiments, animals received 10 µg of EV protein in 100 µL PBS per injection. EV quality was assessed by particle sizing, protein quantification, and immunoblotting for EV-enriched and negative markers.

### Tunable and microfluidic resistive pulse sensing for particle sizing characterization

EV size distribution and concentration were measured by tunable resistive pulse sensing (TRPS) using the qNanoGold platform (Izon Science). Freshly purified EVs were diluted 1:40 in 0.2 um-filtered PBS. For TRPS, LOs were analyzed using an NP2000 nanopore and sEVs using an NP250 nanopore. Membranes were stretched to 47 mm, and voltage was set to 0.04 V for LOs or 0.5 V for sEVs to obtain a stable baseline current of approximately 120 nA. Particle size and concentration were calibrated using Izon calibration particles (CPC2000 diluted 1:1,000 for large EVs and CPC200 diluted 1:100 for sEVs). At least 500 events were recorded per sample at 5 mbar positive pressure.

### Enrichment of circulating LOs from mouse plasma

For circulating EV analysis, blood was collected from mice at endpoint into BD Vacutainer K2 EDTA tubes. Samples were processed within 2 h of collection. Platelet-poor plasma was generated by sequential centrifugation of 500 x g 10 min and 2,000 x g for 20 min. For LO enrichment, approximately 800 uL of plasma was pooled from mice within each experimental group using an equal volume from each animal. Pooled plasma was diluted in 0.2 um-filtered PBS to a final volume of 1 mL and centrifuged at 10,000 x g for 30 min at 4°C. Crude plasma LO pellets were resuspended in 10 µL 0.2 um-filtered PBS and used for downstream direct stochastic optical reconstruction microscopy (dSTORM) analysis of PIGS and Caveolin-1-positive particles.

### Protein quantification and EV dosing

Protein concentration in EV preparations and whole-cell lysates was measured using the Pierce BCA Protein Assay Kit (Thermo Fisher Scientific), according to the manufacturer’s instructions. A BSA standard curve ranging from 20 to 2,000 ug/mL was used, and samples were measured with at least three technical replicates. Buffer compatibility with PBS, RIPA, and iodixanol-containing samples was confirmed before analysis. EV doses for functional assays were normalized to total protein unless otherwise noted.

### Whole-cell/EV lysates and immunoblotting

Whole-cell lysates were prepared from cultured cells between 80-90% confluence. Cell monolayers were washed three times with chilled PBS, scraped, and lysed in RIPA buffer. For EV lysates, after gradient density purification RIPA buffer 10x was added to the EV preparations in PBS to a final 1x concentration for lysis. Lysates were supplemented with Complete protease inhibitor (Roche) and incubated on ice for 20 min followed by a centrifugation at 10,000 x g for 30 min at 4°C. Cleared supernatant was collected and protein concentration determined by BCA assay. EV and whole-cell lysate samples were separated by SDS-PAGE (1-10 µg of protein) and transferred to nitrocellulose or low-fluorescence PVDF membranes. Immunoblotting was performed using primary antibodies including HSPA5 (Cell Signaling Technology, #3177, 1:1,000), cytochrome c (Cell Signaling Technology, #4280, 1:1,000), TSG101 (Santa Cruz Biotechnology, sc-7964, 1:1,000), CD9 (Santa Cruz Biotechnology, sc-13118, 1:1,000), CD81 (Abcam, ab79559, 1:10,000), Caveolin-1 (Santa Cruz Biotechnology, sc-894, 1:10,000), KRT18 (Abcam, ab93741, 1:10,000), STING (D2P2F; Cell Signaling Technology, #13647, 1:1,000), SEPT2 (Sigma, HPA018481, 1:1,000), beta-actin (Sigma, 1:2,500), and GAPDH (Cell Signaling Technology, 1:1,500). HRP-conjugated anti-mouse and anti-rabbit secondary antibodies (Cell Signaling Technology, 1:5,000) and fluorescent StarBright 560 anti-mouse and StarBright 700 anti-rabbit secondary antibodies (Bio-Rad, 1:2,500) were used where appropriate.

### Transmission electron microscopy

Purified EV pellets were fixed with 4% paraformaldehyde and post-fixed in 0.5% osmium tetroxide. Samples were dehydrated through graded ethanol, block stained with 1% uranyl acetate in 50% ethanol for 30 min, embedded in Taab 812 resin, and polymerized overnight at 60°C. Ultrathin sections were imaged using a Hitachi 7100 transmission electron microscope equipped with a Veleta 2000 x 2000 pixel side-mounted CCD camera (Olympus).

### Super-resolution microscopy and dSTORM of EVs

Direct stochastic optical reconstruction microscopy (dSTORM) was performed on an ONI Nanoimager system (Oxford Nanoimaging) using a method developed by our group (Newman et al., *under revision*). Purified sEVs and LOs were diluted 1:100 and 1:40 in PBS, respectively, and 100 µL was loaded into µ-Slide VI 0.5 glass-bottom chambers (Ibidi). EVs were immobilized by centrifugation at 2,000 rpm for 3 min at 4°C, washed twice with PBS, fixed in 4% paraformaldehyde for 10 min, and blocked with 5% BSA for 30 min. Samples were stained for 45 min with the ONI Pan-EV membrane dye, fluorophore-conjugated primary antibodies, or unconjugated primary antibodies followed by fluorescent secondary antibodies. Reagents included RPN2-AF647 (Novus Biologicals; 7 µg/mL), CD81/CD63/CD9-CF568 (ONI EV Profiler Kit; 1:6.25), CD63-AF561 (ONI; 1:25), PIGS (Proteintech; 7 µg/mL), Caveolin-1 (Santa Cruz Biotechnology; 4 µg/mL), dsDNA clone AE-2 (Merck Millipore; 10 µg/mL), and dsRNA clone J2 (Thermo Fisher Scientific; 5 µg/mL). Unconjugated antibodies were detected with anti-rabbit IgG-AF647 (Invitrogen; 5 µg/mL) or anti-mouse IgG-CF568 (Sigma-Aldrich; 5 µg/mL). Secondary-only controls were used to assess nonspecific fluorescence.

For imaging, 100 µL of dSTORM imaging buffer was added to each lane immediately before acquisition. Six fields of view were acquired per lane across consistent lower, middle, and upper regions using an illumination angle of 53°. Raw images were analyzed using ONI CODI software. Localization filtering parameters were defined using secondary-only controls, and Pan-EV membrane dye localizations were used to identify membrane-positive clusters corresponding to individual EVs. Colocalization of protein or nucleic acid markers with membrane-stained EV clusters was quantified using CODI to determine the percentage of single-positive and double-positive EVs. Plasma-derived LO preparations were analyzed using the same workflow to quantify circulating PIGS-positive and Caveolin-1-positive particles.

To distinguish external from intravesicular nucleic acid signals, LO preparations were analyzed under intact, permeabilized, and nuclease-treated conditions. Intact LOs were stained with anti-dsDNA or anti-dsRNA antibodies to detect external nucleic acid signal, whereas permeabilized LOs were stained to detect both external and intravesicular signal. Permeabilization was performed using the EV Profiler Kit permeabilization buffer (ONI). For nuclease treatments, samples were treated on-slide with 1 µL DNase I per lane (Invitrogen TURBO DNase; 2 U/µL) for 30 min at 37°C or 10 µL RNase III per lane (New England Biolabs; 2,000 U/mL), prepared according to the manufacturer’s instructions and incubated for 20 min at 37°C. In intact nuclease-treated LOs, external nucleic acids were digested before staining, whereas nuclease-treated LOs that were subsequently washed and permeabilized were used to assess protected intravesicular nucleic acid signal.

### Vesicle flow cytometry

Single-particle flow cytometry was used to quantify dsRNA-positive particles within the LO-size gate. EV preparations were stained with PE-conjugated J2 antibody to detect dsRNA and FITC-conjugated wheat germ agglutinin (WGA) to identify membrane-bound particles. Samples were incubated for 1 h at room temperature in the presence of 0.1% saponin for permeabilization. Data were acquired on a BD FACSymphony A5 using fluorescence-triggered detection. Size calibration and gating were performed with Rosetta Calibration Beads (Exometry), and analyses were restricted to particles ≥1 um. WGA-positive events were used to define vesicular populations, and dsRNA positivity was quantified within this gate. Buffer-only, single-stained, and fluorescence-minus-one controls were included for gating and compensation. Data were analyzed in FlowJo v11.

### BM-MSC EV treatments and conditioned medium collection

Human or murine BM-MSCs were plated, after 24 h of adherence, serum-free medium was replaced and cells starved for 4 h prior to be exposed to purified LOs, sEVs, or PBS/vehicle control for 24 h. Unless otherwise indicated, *in vitro* EV treatments were performed using 20 ug/mL EV protein. Dose-response experiments used the EV concentrations indicated in the corresponding figures. For transcriptomic and qPCR analyses, RNA was collected at the timepoints indicated in each experiment. For conditioned medium experiments, BM-MSCs were treated with EVs, washed as appropriate, and conditioned medium was collected for cytokine/chemokine measurement, tumor-cell migration, neutrophil migration, and neutrophil polarization assays. All the experiments were performed at least three times, and each experimental group had at least three biological replicates.

### Cytokine, chemokine, and 2’3’-cGAMP detection

Soluble cytokines and chemokines released by EV-treated BM-MSCs were measured in conditioned medium using the Human Cytokine/Chemokine Panel A 48-Plex Discovery Assay® Array (HD48A; Eve Technologies), according to the manufacturer’s protocol. Briefly, conditioned media were collected from BM-MSC cultures following treatment with LOs, sEVs, or vehicle control, clarified to remove cellular debris, and stored at −80°C until analysis. Samples were submitted to Eve Technologies for multiplex quantification of cytokines and chemokines. Analyte concentrations were calculated from standard curves generated for each target and reported as concentration per volume of conditioned medium.

2’-3’-cyclic GMP-AMP (2’-3’-cGAMP) was quantified using the 2’-3’-cyclic GAMP ELISA Kit (Invitrogen), according to the manufacturer’s instructions. Donor cancer cells, purified LOs, and sEVs were processed for cGAMP detection using 20 µg of total protein per sample as input for normalization. Absorbance was measured using a microplate reader at the wavelength recommended by the manufacturer, and 2’-3’-cGAMP concentrations were calculated from the kit standard curve. Values below the assay detection limit were handled as zero for downstream analysis.

### RNA extraction, RT-qPCR, and primer information

Total RNA was extracted from cells using the Quick-RNA Miniprep Kit (Zymo Research), according to the manufacturer’s instructions. Gene expression was analyzed by one-step reverse transcription quantitative PCR (RT-qPCR) using the iTaq Universal One-Step RT-qPCR Kit (Bio-Rad) on a QuantStudio 5 Real-Time PCR System. Briefly, 40 ng of total purified RNA was added directly to one-step RT-qPCR reactions containing reverse transcriptase, universal reaction mix, and gene-specific primer pairs. Reverse transcription and quantitative PCR amplification were performed in the same reaction according to the manufacturer’s recommended cycling conditions. Relative gene expression was normalized to housekeeping genes and calculated using the ΔΔCt method.

The following primer pairs were used for inflammatory, interferon-stimulated, nucleic acid-sensing, and immunosuppressive genes. Human primers included: CCL2, forward 5’-AGAATCACCAGCAGCAAGTGTCC-3’ and reverse 5’-TCCTGAACCCACTTCTGCTTGG-3’; CCL5, forward 5’-CCTGCTGCTTTGCCTACATTGC-3’ and reverse 5’-ACACACTTGGCGGTTCTTTCGG-3’; CXCL1, forward 5’-AGCTTGCCTCAATCCTGCATCC-3’ and reverse 5’-TCCTTCAGGAACAGCCACCAGT-3’; CXCL8, forward 5’-GAGAGTGATTGAGAGTGGACCAC-3’ and reverse 5’-CACAACCCTCTGCACCCAGTTT-3’; IL6, forward 5’-AGACAGCCACTCACCTCTTCAG-3’ and reverse 5’-TTCTGCCAGTGCCTCTTTGCTG-3’; CXCL10, forward 5’-GGTGAGAAGAGATGTCTGAATCC-3’ and reverse 5’-GTCCATCCTTGGAAGCACTGCA-3’; CXCL11, forward 5’-AAGGACAACGATGCCTAAATCCC-3’ and reverse 5’-CAGATGCCCTTTTCCAGGACTTC-3’; RSAD2, forward 5’-CCAGTGCAACTACAAATGCGGC-3’ and reverse 5’-CGGTCTTGAAGAAATGGCTCTCC-3’; ARG1, forward 5’-TCATCTGGGTGGATGCTCACAC-3’ and reverse 5’-GAGAATCCTGGCACATCGGGAA-3’; IL10, forward 5’-TCTCCGAGATGCCTTCAGCAGA-3’ and reverse 5’-TCAGACAAGGCTTGGCAACCCA-3’; and TNF, forward 5’-CTCTTCTGCCTGCTGCACTTTG-3’ and reverse 5’-ATGGGCTACAGGCTTGTCACTC-3’. Murine primers included: Ccl5, forward 5’-CCTGCTGCTTTGCCTACCTCTC-3’ and reverse 5’-ACACACTTGGCGGTTCCTTCGA-3’; Cxcl10, forward 5’-ATCATCCCTGCGAGCCTATCCT-3’ and reverse 5’-GACCTTTTTTGGCTAAACGCTTTC-3’; Rsad2, forward 5’-GGAAGGTTTTCCAGTGCCTCCT-3’ and reverse 5’-ACAGGACACCTCTTTGTGACGC-3’; Irf3, forward 5’-CGGAAAGAAGTGTTGCGGTTAGC-3’ and reverse 5’-CAGGCTGCTTTTGCCATTGGTG-3’; Irf7, forward 5’-CCTCTGCTTTCTAGTGATGCCG-3’ and reverse 5’-CGTAAACACGGTCTTGCTCCTG-3’; Ifna1, forward 5’-GGATGTGACCTTCCTCAGACTC-3’ and reverse 5’-ACCTTCTCCTGCGGGAATCCAA-3’; and Ddx58/Rig-I, forward 5’-AGCCAAGGATGTCTCCGAGGAA-3’ and reverse 5’-ACACTGAGCACGCTTTGTGGAC-3’. Housekeeping gene primers included human GAPDH, forward 5’-GTCTCCTCTGACTTCAACAGCG-3’ and reverse 5’-ACCACCCTGTTGCTGTAGCCAA-3’; mouse Gapdh, forward 5’-CATCACTGCCACCCAGAAGACTG-3’ and reverse 5’-ATGCCAGTGAGCTTCCCGTTCAG-3’; human ACTB, forward 5’-CACCATTGGCAATGAGCGGTTC-3’ and reverse 5’-AGGTCTTTGCGGATGTCCACGT-3’; and mouse Actb, forward 5’-CATTGCTGACAGGATGCAGAAGG-3’ and reverse 5’-TGCTGGAAGGTGGACAGTGAGG-3’. The housekeeping gene used for each experiment was selected according to the species of the biological sample.

### RNA sequencing and transcriptomic analysis of EV-treated BM-MSCs

RNA-seq was performed using three biological replicates per group. RNA was extracted from experimental samples, treated with TURBO DNase (Thermo Fisher Scientific, AM2238), and cleaned using the Zymo RNA Clean and Concentrator kit (Zymo Research, R1014). RNA concentration and quality were assessed using RiboGreen reagent (Thermo Fisher Scientific, R11490) and Agilent TapeStation RNA ScreenTape reagents (5067-5579). 2.5 ng of RNA was used for library preparation with the SMARTer Pico V2 kit (Takara, 634412). Libraries were quantified using KAPA Library Quantification Kits (Roche, KK4824) and sequenced as single-end reads on an Illumina NovaSeq 6000.

Transcripts were quantified using Salmon, and differential expression analysis was performed with DESeq2. Heatmaps were generated using the ComplexHeatmap R/Bioconductor package on scaled log-expression values (z-scores), with Euclidean distance and Ward linkage implemented with the fastcluster package. For selected heatmaps, genes with absolute log2 fold change >1 were ranked by P value and the top genes were plotted. Gene Set Enrichment Analysis (GSEA) was performed on genes ranked by fold change using clusterProfiler and MSigDB gene-set collections. Significantly enriched gene sets were reported using Benjamini-Hochberg adjusted P values <0.05.

### Innate immune reporter assays and pathway perturbation

Innate immune activation by LOs was evaluated using a BM-MSC reporter-cell assay based on secreted embryonic alkaline phosphatase (SEAP) activity. BM-MSCs were transfected with the pNiFty2-56K-SEAP plasmid (InvivoGen) using Lipofectamine™ Stem Transfection Reagent. The plasmid was first propagated in Mix & Go! DH5α competent cells, and bacterial clones were selected with zeocin at 25 µg/mL. Following transfection, Zeocin™-resistant BM-MSCs were selected using zeocin at 100 µg/mL to generate reporter BM-MSCs. Reporter activity was measured using QUANTI-Blue™ SEAP Detection Reagent (InvivoGen), according to the manufacturer’s instructions. For reporter assays, BM-MSC reporter cells were plated in 96-well plates at 5×10^3^ cells per well and allowed to adhere for 24 h. The medium was then replaced with serum-free medium before treatment. Cells were exposed to LOs at 20 µg/mL. Poly(I:C) and poly(dA:dT) complexed with transfection reagent LyoVec (Invivogen) were used as positive controls for nucleic acid-induced innate immune activation and were added at the same time as LOs at final concentrations of 1 µg/mL and 10 µg/mL, respectively. To assess STING-dependent signaling, reporter BM-MSCs were pretreated with the STING inhibitor H-151 (MedChemExpress) at a final concentration of 15 µM. H-151 was added 4 h before LO, poly(I:C), or poly(dA:dT) exposure and was maintained in the culture medium throughout LO or agonist stimulation. Thus, STING signaling was inhibited before and during exposure to EV-associated cargo. DMSO was used as the vehicle control at the same volume used in inhibitor-treated groups. A toxicity curve was performed to select inhibitor conditions that did not induce detectable cell death.

To evaluate whether endosomal acidification or degradation influenced LO-induced inflammatory signaling, BM-MSCs were treated with Bafilomycin A1 (MedChemExpress) at a final concentration of 100 nM. Bafilomycin A1 was added 2 h before LO exposure and maintained in the medium during LO stimulation (24 h). DMSO was used as the vehicle control at the same volume used in the Bafilomycin A1-treated group. For both studies, LO-induced inflammatory responses following pathway perturbation were measured by either SEAP reporter activity and/or downstream gene-expression readouts, as described in the RT-qPCR section.

### Generation of NanoLuc/GFP reporter 4T1.2 cells

NanoLuciferase/GFP reporter-expressing 4T1.2 cells were generated by lentiviral transduction. Lentiviral particles were produced in HEK293T cells by co-transfection of pLenti-PalmGRET (Addgene #158221, gift from Dr Charles P. Lai) with packaging plasmids psPAX2 (Addgene #12260) and pMD2.G (Addgene #12259) using TurboFectin 8.0. Viral supernatants were collected at 48 and 72 h post-transfection, filtered through a 0.45 µm filter, and stored at −80 C. 4T1.2 cells were transduced with lentiviral supernatant in the presence of 8 ug/mL polybrene overnight. GFP-positive cells were enriched by bulk FACS sorting, and the top 20% brightest GFP-positive cells were collected, expanded, and selected with puromycin (5 ug/mL).

### Immunofluorescence microscopy of cells

Cellular immunofluorescence was used to visualize bleb formation in the plasma membrane and dsRNA accumulation/localization. Cells seeded in round cover class, after 24 h cells were fixed and permeabilized using BD Cytofix/Cytoperm Fixation/Permeabilization Solution Kit, and stained with dsRNA J2 antibody (Jena Bioscience, rabbit-raised, 1:100 for cell imaging), rhodamine-phalloidin (Cytoskeleton Inc., #PHDR1, 100 nM), WGA-AF488 or WGA-AF555 (Invitrogen, #W11261/#W32464, 1 ug/mL), and DAPI (Invitrogen, #62248, 1 ug/mL) across different experiments. Cells were visualized in Leica Stellaris 8-STED Super-resolution Confocal Microscope.

### Tumor-cell migration and dynamic bone-on-chip assay

Cancer cell and neutrophil migration toward conditioned medium from EV-treated BM-MSCs was evaluated using transwell migration assays. BM-MSCs were treated with LOs, sEVs, or vehicle control, and conditioned medium was collected as described above. PC3 cells or freshly isolated human neutrophils were seeded into the upper chamber of transwell inserts containing 8.0 µm pore-size PET membranes. Conditioned medium from control or EV-treated BM-MSCs was placed in the lower chamber as the chemoattractant. After migration, non-migrated cells were removed from the upper surface of the membrane, and migrated cells on the lower surface were fixed, stained with crystal violet, and quantified by counting the number of migrated cells per insert or imaging field.

For dynamic 3D bone-on-a-chip assays, we used a previously described PDMS microfluidic platform that supports BM-MSC culture under continuous perfusion and microscopic imaging^[30]^. Chips were coated with laminin, and BM-MSCs were seeded at 1×10^6^ cells/mL in 50 µL. After 3-6 h, non-adherent cells were removed by flushing, and flow was initiated 24 h later at 30 µL/h using a syringe pump. LOs or vehicle were then circulated through the chip to condition the stromal compartment. PC3 cells were subsequently introduced into the circulating medium, and their recruitment and attachment to the BM-MSC-seeded matrix were quantified by imaging as the number of matrix-associated cells per field or region of interest. This system enabled assessment of LO-mediated stromal education under physiologically relevant flow, matrix, and shear conditions.

### CRISPR-Cas9 knockout of STING and SEPT2

CRISPR-Cas9 knockout of human STING/TMEM173 in BM-MSCs and mouse Sept2 in 4T1.2 cells was performed by ribonucleoprotein (RNP) delivery using the Alt-R CRISPR-Cas9 system (Integrated DNA Technologies) and Lipofectamine CRISPRMAX (Thermo Fisher Scientific), according to the manufacturer’s recommendations. Gene-specific crRNAs were annealed with tracrRNA (1:1 molar ratio) in Nuclease Free Duplex Buffer (Integrated DNA Technologies) to generate duplex guide RNAs, and incubated at 95 °C for 5 min followed by continuous ramp down temperature step of −1 °C/min. Annealed gRNAs were complexed with RFP-SpCas9 recombinant protein before transfection (1.2:1 molar ratio, respectively). The TMEM173-targeting crRNA sequence was 5-GCUGGGACUGCUGUUAAACGGUUUUAGAGCUAUGCU-3. The Sept2-targeting crRNA sequence was 5-UAGUGGACACUCCCGGCUACGUUUUAGAGCUAUGCU-3. A scrambled sequence not targeting anything sequence on the human or murine genome was used as a negative control 5-CGUUAAUCGCGUAUAAUACGGUUUUAGAGCUAUGCU-3. RFP-positive cells were isolated by flow sorting approximately 24 h after transfection. STING knockout BM-MSCs were enriched by bulk sorting, whereas 4T1.2 Sept2-edited cells were sorted as single cells to generate monoclonal populations. Knockout was validated by western blotting.

### Flow cytometry and high-dimensional analysis

Flow cytometry was used to assess neutrophil polarization *in vitro* and to profile bone marrow immune populations *in vivo*. For human neutrophil polarization assays, freshly isolated human neutrophils were exposed for 1–6 h to conditioned medium collected from BM-MSCs treated with LOs, sEVs, or vehicle control. After stimulation, cells were stained with LOX-1-FITC (Thermo Fisher Scientific, 1:100), CD66b-PE (Thermo Fisher Scientific, 1:100), and CD62L-AF647 (1:50) to evaluate acquisition of a PMN-MDSC-like phenotype.

For *in vivo* immune profiling, bone marrow cells were harvested from experimental mice, filtered into single-cell suspensions, subjected to red blood cell lysis with ACK lysis buffer, counted, and stained for spectral flow cytometry. Cells were processed using the BD Cytofix/Cytoperm kit according to the manufacturer’s recommendations for fixation and permeabilization/blocking prior to acquisition. The staining panel included CD45-PE-Cy7 (BD, 1:100), CD3-PE-Cy5 (BD, 1:100), CD11b-BB700 (BD, 1:200), CD11c-BV421 (BD, 1:200), F4/80-BUV496 (BD, 1:200), Ly6C-FITC (BD, 1:100), Ly6G-BV563 (BD, 1:200), CD14-APC (BD, 1:200), CD16/32-BUV737 (BD, 1:200), MHC-II-BV605 (BD, 1:200), B220-PerCP (BD, 1:200), CD177-AF647 (BD, 1:200), PD-L1-BV711 (BD, 1:100), CD49d-BV786 (BD, 1:200), CD86-BV510 (BD, 1:200), CXCR2-BUV661 (BD, 1:100), CXCR4-PE (BD, 1:100), TREM-1-BUV395 (BD, 1:200), CD64-BV650 (BD, 1:200), CD62L-BUV805 (BD, 1:200), Siglec-F-BV750 (BD, 1:200), and BD Horizon Fixable Viability Stain 780 (1:1,000) ^[31–33]^. Samples were acquired on a Sony ID7000 spectral flow cytometer. Autofluorescence was defined using unstained controls and removed during spectral unmixing. Single-stained reference controls for each fluorophore were used to generate spectral reference signatures and perform unmixing. A total of 15,000 events were acquired. Gating was performed in FlowJo v10 after exclusion of debris, doublets, and dead cells, followed by biexponential transformation of fluorescence parameters. Neutrophils and PMN-MDSC-like populations were identified using a sequential gating strategy that included live cells, CD45⁺ leukocytes, CD11b⁺ myeloid cells, Ly6G⁺ neutrophils, and additional phenotypic markers including Ly6C, CD14, PD-L1, CXCR2, MHC-II, CD62L, CD177, and related myeloid markers^[33]^. For high-dimensional analysis, live cells from five mice were concatenated and exported as a single FCS file. Dimensionality reduction visualization was performed using the Uniform Manifold Approximation and Projection (UMAP) algorithm. FlowSOM clustering^[34]^ was then applied and overlaid onto the UMAP projection to objectively identify phenotypically distinct populations based on marker-expression patterns. Metaclusters with similar expression profiles were manually merged when appropriate and annotated according to the expression of defining markers.

### Scaffold and intratibial neutrophil recruitment assays

For *in vivo* neutrophil recruitment assays, mouse bone marrow cells or MSCs were seeded either into 24-well plates or into 3D scaffold disks. Scaffolds consisted of reticulated polycarbonate polyurethane urea matrix disks (5 mm x 2 mm; CS1-0502-25, Biomerix Corp/DSM Biomedical). Cells were seeded at 1 x 10^5^ cells per scaffold or per well and allowed to adhere for at least 24 h^[35]^. Cells were then treated with EVs for 24 h. EV-conditioned scaffolds were implanted subcutaneously into C57BL/6 mice flanks, whereas EV-conditioned cells from wells were injected directly into tibiae. Scaffolds and bone marrow cells were harvested 48 h later, dissociated, and analyzed by flow cytometry for neutrophil recruitment (CD45^+^ Cd11b^+^ Ly6C^neg^ Ly6G^+^).

### SEPT2 knockout tumor model and LO rescue

To test whether tumor-derived LO production is required for bone marrow PMN-MDSC accumulation, female 6-8-week-old BALB/c mice were orthotopically implanted in the lower right mammary fat pad with 2×10^5^ scramble control or Sept2 knockout 4T1.2 cells (n=5 mice/group). Tumor growth was monitored during the 2-week experimental period. In rescue cohorts, mice bearing Sept2 knockout or scramble control tumors received systemic LO administration concomitantly with primary tumor growth, consisting of four retro-orbital injections of 10 ug purified syngeneic LO protein in 100 uL PBS over the 2-week period, as described in the Orthotopic 4T1.2 tumor education and metastasis model section. At endpoint, plasma and bone marrow were collected for circulating LO dSTORM and spectral flow cytometry analysis.

### Tumor-bearing versus naive bone marrow immune profiling

To evaluate bone marrow immune remodeling during tumor progression, female 6-8-week-old BALB/c mice were orthotopically implanted with 2×10^5^ nLuc-GFP 4T1.2 cells into the lower right mammary fat pad and compared with naive controls. Tumors were followed for 4 weeks (28 days), after which bone marrow and plasma were collected. Circulating plasma EVs were analyzed by dSTORM for PIGS-positive particles, and bone marrow cells were analyzed by high-dimensional spectral flow cytometry. GFP-positive tumor cells in the bone marrow were assessed to determine whether immune remodeling occurred before overt metastatic colonization.

### IVIS imaging and metastatic burden quantification

Bioluminescence imaging was used to monitor metastatic progression in mice injected with GFP/NanoLuc reporter 4T1.2 cells. Animals were anesthetized before imaging and imaged longitudinally using an IVIS system. Nano-Glo Fluorofurimazine *In vivo* Substrate (Promega) was injected intraperitoneally (0.5 µM), and Signal intensity was quantified as total bioluminescence flux within defined regions of interest, and focal lesions were counted using predefined thresholds based on autofluorescence signal.

### Statistical analysis

Statistical analyses were performed using GraphPad Prism 9 and/or R 4.6. Data are presented as mean ± Standard Error Mean or as otherwise indicated in the figure legends. Two-group comparisons were performed using parametric or nonparametric tests as appropriate, and multiple-group comparisons were analyzed using ANOVA or nonparametric alternatives with post hoc correction. Longitudinal tumor growth or IVIS measurements were analyzed using repeated-measures or mixed-effects models when appropriate. Correlations were assessed using Pearson or Spearman tests based on gaussian distribution characteristics of the sample. RNA-seq analyses used Benjamini-Hochberg adjustment for multiple testing, with adjusted P <0.05 considered significant unless otherwise specified. The definition of n corresponds to mice, biological replicates, independent EV preparations, or human donors as specified in each figure legend.

## RESULTS

### LOs enhance tumor cell migration and metastatic progression

Given the emerging role of tumor-derived EVs in pre-metastatic niche formation and systemic tumor-host communication, we decided to investigate whether LOs and sEVs differ in their capacity to influence metastatic progression *in vivo*. To address this question, we employed a syngeneic breast cancer model (4T1.2 cells) with bone metastatic propensity, recapitulating spontaneous bone metastasis in breast cancer patients^[36,37]^. Using a two-phase “education” approach, immunocompetent BALB/c mice bearing orthotopic mammary fat pad tumors received systemic retro-orbital administration of either LOs or sEVs derived from 4T1.2 cell line throughout primary tumor progression (**Fig. 1A**).

**Figure 1.**
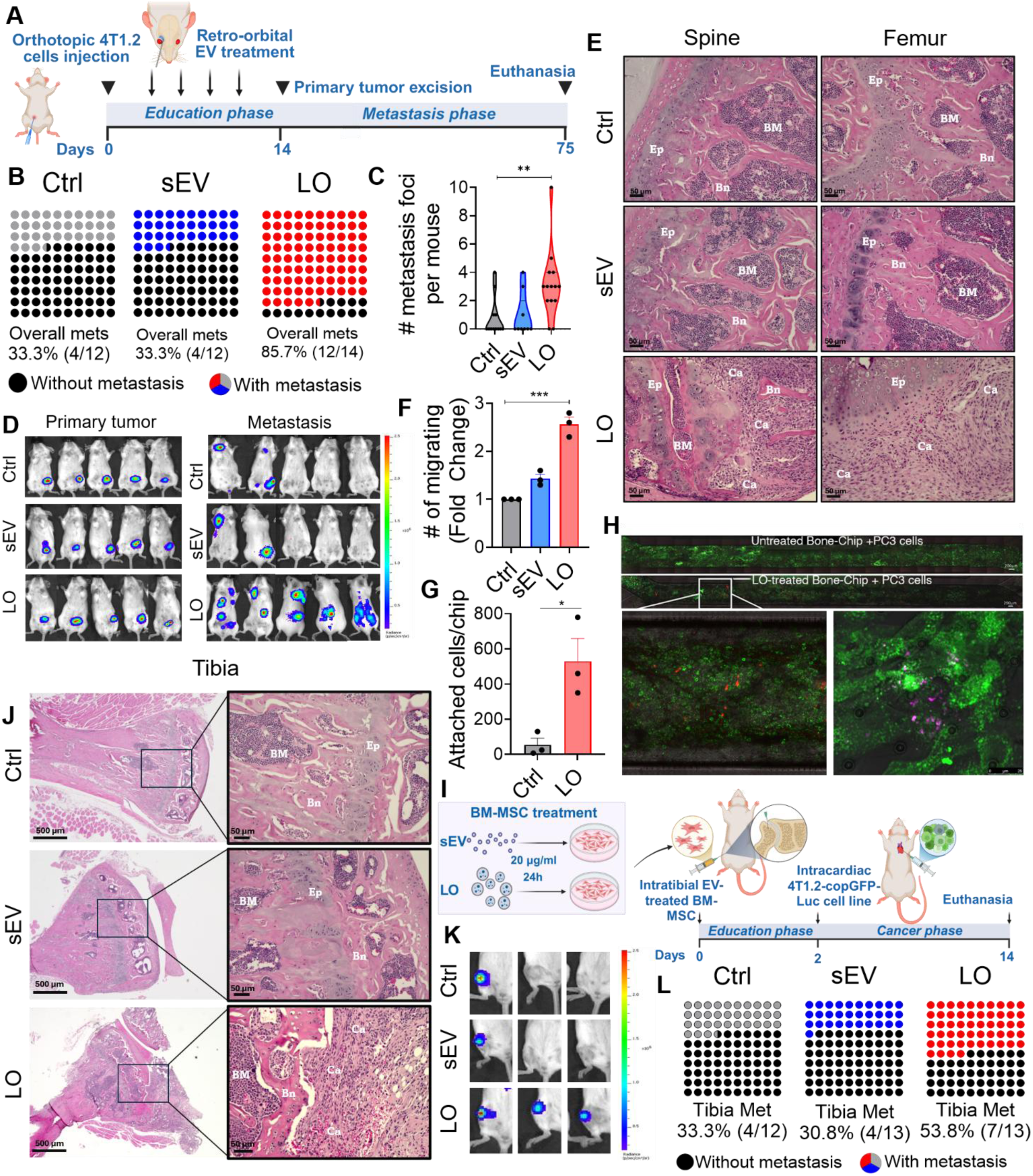
Large oncosomes exhibit enhanced pro-metastatic activity. **A**, Schematic of the metastasis-education model using orthotopic implantation of 1×10^5^ 4T1.2-Luc/GFP cells in BALB/c mice followed by intravenous administration of 10 µg/injection of LO, sEV or vehicle (education phase) derived from 4T1.2 cells. Primary tumors were excised and metastasis development evaluated for additional 60 days (metastasis phase), n ≥ 14 mice per group. **B**, Waffle plots showing metastatic incidence in the Ctrl-, sEV- and LO-educated groups. **C**, Quantification of metastatic foci in each experimental group. **D**, Representative IVIS images of mice from each group at the end of the end of education phase (left) and at endpoint (60 days after primary tumor excision; right). **E,** Hematoxylin and Eosin (H&E) staining of spine and femur sections. Scale bar 50 µm. **F**, Transwell recruitment assay in which PC3 cells migrate toward conditioned medium from BM-MSCs treated with PC3-derived LO, sEV or Ctrl. **G-H**, quantification and imaging of bone-on-a-chip assay showing RFP-PC3 (red) migration toward GFP-MSCs (green) pre-exposed to LO. Internalized RFP-PC3-derived LO are shown in the left panel (magenta), and quantification of total number of cancer cells attaching to the BM-MSC containing matrix shown in the right. Scale bar 200 and 25 µm. **I**, Schematic of the MSC-priming bone metastasis model: BALB/c BM-MSCs pretreated for 24 h with 4T1.2-derived LO, sEV or vehicle were injected (2×10^4^) into the tibia of Balb/c. After 48 h mice received intracardiac injection of 1×10^5^ 4T1.2-Luc/GFP cells and tibia colonization was analyzed after 14 days. n ≥ 12 mice per group. **J,** H&E staining of tibia sections **K**, Representative IVIS images of the knee region at endpoint. **L**, Waffle plots showing metastatic incidence in tibias pre-injected with MSCs treated with LO, sEV, and Ctrl. Ep – epiphysis; BM – bone marrow; Bn – bone; Ca – Cancer cells.

LO-treated mice showed a marked increase in metastatic incidence, with metastases detected in 85.7% of animals compared with 33.3% in both control and sEV-treated groups (**Fig. 1B**). Consistent with this, LO education significantly increased the number of metastatic foci per mouse, indicating that LOs promote not only metastatic occurrence but also overall metastatic burden (**Fig. 1C**). Importantly, this result was not explained by differences in primary tumor growth, as IVIS imaging showed comparable primary tumor signal across groups, while revealing broader metastatic dissemination in LO-treated animals after primary tumor excision (**Fig. 1D**). Histological analysis further supported these findings. Spine and femur sections from control and sEV-treated mice showed largely preserved bone architecture, including defined epiphyseal regions, intact bone structures, and normal-appearing bone marrow. In contrast, bones from LO-treated mice displayed disrupted architecture, altered epiphyseal organization, reduced bone area consistent with the osteolytic behavior of this model, and extensive replacement of bone marrow by tumor cells (**Fig. 1E**). Together, these data indicate that systemic exposure to LOs during primary tumor progression enhances metastatic colonization and bone involvement without increasing primary tumor growth.

The ability of LOs to enhance metastatic dissemination prompted us to examine their impact on stromal populations that regulate metastatic niche formation. We investigated whether LOs contribute to stromal education through the modulation of BM-MSCs, which are known to play a key role in establishing a supportive niche for metastatic tumor cells^[38–40]^. LO-treated BM-MSCs significantly enhanced prostate cancer PC3 cells migration compared to sEV in an *in vitro* transwell assay (**Fig. 1F**). This was further confirmed using a dynamic 3D bone-on-a-chip, where LO-primed BM-MSCs recruited substantially more prostate cancer cells than naïve BM-MSCs (**Fig. 1G-H**).

To determine whether LO-conditioned stromal cells are sufficient to enhance metastatic colonization *in vivo*, BALB/c-derived BM-MSCs were pre-treated with 4T1.2-derived LOs or sEVs and injected intratibially into syngeneic BALB/c mice prior to intracardiac inoculation of 4T1.2 tumor cells (**Fig. 1I**). This approach allowed us to test whether local stromal education by tumor-derived vesicles creates a more permissive bone microenvironment for subsequent metastatic seeding. Histological analysis of the injected tibias showed preserved bone architecture in control and sEV-primed groups, whereas tibias receiving LO-conditioned BM-MSCs displayed overt tumor infiltration and disruption of the bone marrow compartment (**Fig. 1J**). Consistent with these findings, IVIS imaging revealed focal metastatic signal in the injected tibial region of LO-primed animals (**Fig. 1K**). Quantification across the cohort showed that tibial metastases developed in 53.8% of mice receiving LO-conditioned BM-MSCs, compared with 33.3% and 30.8% of mice in the control and sEV-conditioned BM-MSC groups, respectively (**Fig. 1L**). Together, these data suggest that LO-mediated reprogramming of BM-MSCs enhances the ability of the local bone microenvironment to support metastatic colonization.

### LOs are structurally distinct EVs and trigger a robust interferon-driven inflammatory program in BM-MSCs

Having defined their foundational functional properties, we next sought to investigate the molecular pathways activated in BM-MSCs upon LO exposure. To characterize the physical and molecular features of LOs, we first performed super-resolution microscopy on LOs and sEVs isolated from bone metastatic prostate cancer PC3 cells. sEV identity was confirmed by positivity to a tetraspanin trio (CD9, CD63, and CD81) and LO identity by positivity to CD63 and RPN2^[9]^. Individual LOs displayed micrometer-scale dimensions, with particles >1 μm in diameter (**Fig. 2A-B and S1A**).

**Figure 2.**
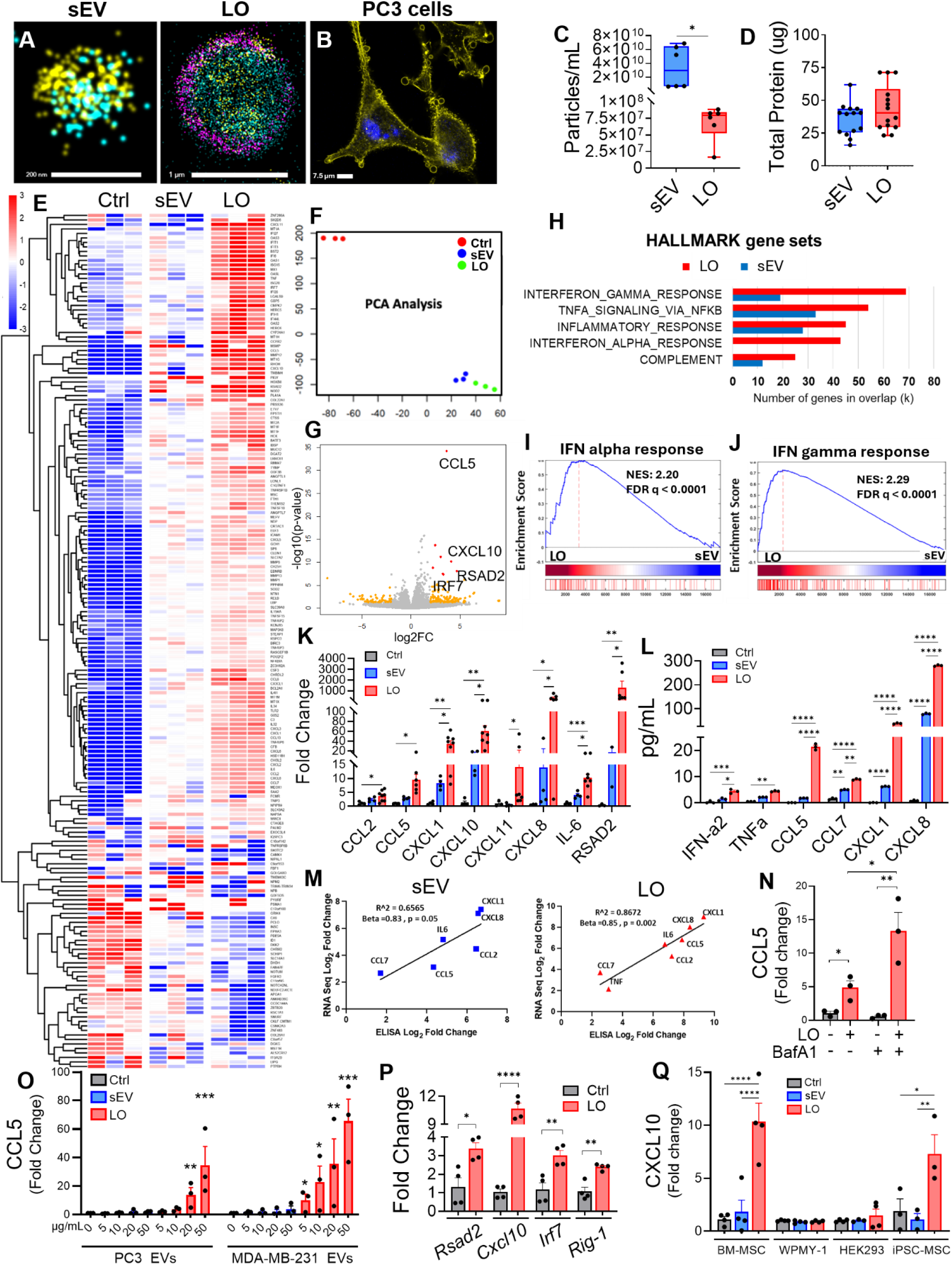
Large oncosomes induce a more robust inflammatory transcriptional program in human bone marrow mesenchymal stem cells than S-EVs. **A**, super-resolution direct Stochastic Optical Reconstruction Microscopy (dSTORM) of PC3-derived small EVs (sEV) and large oncosomes (LO) labeled with a pan–extracellular vesicle membrane dye (cyan) and population-specific markers. sEV were identified using tetraspanins CD9, CD63 and CD81 (yellow), whereas LOs were marked by CD63 (yellow) and RPN2 (magenta). Scale bar, 200 nm for sEV and 1 µm for LO. **B**, Confocal microscopy imaging of PC3 cells stained with phalloidin-rhodamine (yellow) and DAPI (blue) showing bebbling formation. **C,** Tunable-resistive pulse sensing (TRPS) particle concentration measurements of PC3-derived LO and sEVs. **D**, Total protein (µg) abundance of PC3-derived LO and sEV preparations. Each dot represents one independent experiment. **E**, RNA sequencing (RNA-seq) performed on human BM-MSCs treated for 24 h with PC3-derived large oncosomes (LO), sEV, or vehicle (PBS). Heatmap summarizing differentially expressed genes across RNA-seq samples from LO-, sEV- and vehicle-treated human BM-MSCs (n = 3 biological replicates per group). RNA-seq differential expression analysis was performed using log2 fold-change ranking with multiple-testing correction as described in the Methods. **F**, Principal component analysis (PCA) of RNA-seq samples showing clustering by treatment condition. **G**, Volcano plot showing differential gene expression in LO-treated versus vehicle-treated human BM-MSCs. Points are colored according to log2 fold-change and adjusted q-value thresholds, Log2FC > 1 or < −1; q < 0.05. **H**, Gene set enrichment analysis (GSEA) of MSigDB Hallmark gene sets (v7.0), showing the top enriched pathways in LO-treated versus vehicle-treated human BM-MSCs, with corresponding overlap in sEV-treated versus vehicle-treated cells displayed for comparison. Bars indicate the number of overlapping genes. **I-J**, GSEA enrichment plots for interferon alpha and gamma response in LO-treated versus sEV-treated BM-MSCs. **K**, RT–qPCR analysis of inflammatory gene expression in human BM-MSCs treated with LO, sEV or vehicle. Each dot represents one independent experiment. **L**, ELISA quantification of cytokines in supernatants from human BM-MSCs treated with LO, sEV or vehicle. **M**, Correlation between RNA-seq fold changes and cytokine protein levels measured by ELISA in human BM-MSCs treated with sEV or LO. **N**, RT–qPCR analysis of CCL5 expression in human BM-MSCs treated with PC3-derived LO following pre-treatment with bafilomycin A1 (100 nM, 2 h). All RT– qPCR expression experiments were normalized to ACTB. **O**, RT–qPCR analysis of CCL5 expression in human BM-MSCs treated with increasing concentrations of LO or sEV derived from PC3 or MDA-MB-231 cells. **P**, RT–qPCR analysis of *Rsad2, Cxcl10, Irf7* and *Rig-1 (Ddx58*) expression in mouse BM-MSCs treated with 4T1.2-derived LO. Expression was normalized to *Actb*. **Q**, CXCL10 expression analysis of BM-MSC, WPMY-1 (fibroblast), HEK293 (epithelial), and iPSC-MSC cells treated with either sEV or LO for 24 h. For all in vitro experiments, data represents at least three independent experiments with at least three biological and three technical replicates. Statistical analyses are described in the Methods.

Tunable resistive pulse sensing (TRPS) highlighted a modal distribution of LO diameter of 1,744 nm in contrast with a modal distribution of sEV diameter of 120.8 nm (**Fig. 2C and S1A**). Transmission electron microscopy (TEM) confirmed the presence of intact, membrane-enclosed vesicles in both the LO and sEV preparations (**Fig. S1B**). Western blotting further demonstrated enrichment of the LO-associated proteins HSP60 and HSPA5 in LOs, whereas the canonical sEV markers TSG101, CD9, and CD81 were enriched in sEVs, with no detectable cellular contaminants in either EV fractions (**Fig. S1C**). Although sEVs were significantly more abundant than LOs, the total protein content was comparable between the two fractions (**Fig. 2C-D**), indicating a higher protein amount per single LO than per single sEV, proportionally to their relative volumes. A similar pattern was observed in mouse-derived bone metastatic 4T1.2 breast cancer cells (**Fig. S1D-H**).

To define the transcriptional responses induced by EVs within the bone mesenchymal niche, RNA-seq on primary human BM-MSCs exposed to prostate cancer cell-derived LOs or sEVs revealed substantially broader transcriptional reprogramming following LO treatment, including a large set of uniquely regulated transcripts, whereas comparatively few genes were altered specifically by sEVs (**Fig. 2E-F** and **Fig. S2A-B**).

LO-treated BM-MSCs displayed marked upregulation of inflammatory and interferon-stimulated genes (ISG), including CCL5, CXCL10, RSAD2, IRF7, ISG15, IFIT family members, and IL6 (**Fig. 2E** and **G**). Hallmark GSEA of the full ranked gene lists identified IFNγ response, TNFα signaling via NF-κB, and inflammatory response among the programs induced by both EV populations. In contrast, the IFNα response was selectively enriched in LO-treated cells and ranked among the most strongly upregulated Hallmark pathways only in the LO condition (**Fig. 2H-J**).

To resolve how these transcriptional programs were distributed across distinct gene modules, we next performed unsupervised clustering of differentially expressed genes followed by cluster-specific Hallmark GSEA. Confirming the previous analysis, genes induced by both LOs and sEVs formed a shared inflammatory cluster. By contrast, genes preferentially induced by LOs formed a distinct cluster in which the IFNα response was the top enriched pathway (**Fig. S2C-D**). Thus, although LOs and sEVs derived from the same cancer cell model, they elicit partially overlapping inflammatory responses; in particular, LOs additionally induce a prominent type I interferon-associated antiviral program, identifying them as distinct modulators of the bone marrow stromal niche.

RT-qPCR confirmed marked upregulation of interferon-stimulated genes CCL2, CCL5, CXCL1, CXCL8, IL-6, CXCL10, CXCL11, and RSAD2 in LO-treated BM-MSCs significantly more than in cells exposed to sEVs (**Fig. 2K**). The changes in transcript abundance identified by RNA-seq in BM-MSCs were strongly reflected at the protein level in their conditioned medium (**Fig. 2L**), with ELISA measurements showing a high degree of concordance with the corresponding gene expression changes (**Fig. 2M**). LO treatment significantly induced CCL5 expression in BM-MSCs, an induction that was markedly enhanced by Bafilomycin A1. Because Bafilomycin A1 alone did not increase CCL5 expression, these data indicate that inhibition of endosomal acidification and lysosomal degradation potentiate LO-mediated inflammatory activation. This suggests that endolysosomal processing may regulate the availability of LO-associated cargo for innate immune sensing in BM-MSCs (**Fig. 2N**).

Further experiments revealed that this response is dose-dependent, regardless of the cancer cell type, for both prostate and breast cancer-derived EVs (**Fig. 2O** and **S2F-G**). The response is also conserved among different species, since murine 4T1.2-derived LOs similarly induced Interferon Stimulated Genes (ISG) such as RSAD2, CXCL10, IRF7, and RIG-I in murine BM-MSCs (**Fig. 2P**). LO treatment induced a robust increase in CXCL10 expression in BM-MSCs. In contrast, WPMY-1 and HEK293 cells remained largely unresponsive to LO exposure, suggesting that LO-induced response might not be a universal response across cell types. Notably, iPSC-derived MSCs also exhibited a significant response to LOs, supporting the idea that mesenchymal cells in the stroma are particularly sensitive to LO-mediated inflammatory reprogramming (**Fig. 2Q**). Together, these findings establish LOs as the EV population that induces a robust, conserved, and dose-dependent interferon-driven inflammatory program in MSCs.

### LO-mediated inflammatory reprogramming is driven by convergent cytosolic DNA- and RNA-sensing pathways

To identify the upstream pathways underlying LO-induced inflammatory response in BM-MSCs, we focused on transcripts selectively upregulated by LOs, which formed a distinct expression profile (**Fig. 3A**), enriched for cytosolic nucleic acid sensing and STING-mediated antiviral pathways (**Fig. 3B**). The result was confirmed by KEGG analysis, which revealed a specific LO-induced enrichment of a cytosolic DNA-sensing pathway (**Fig. 3C**), in line with the presence of both internal and external DNA in LOs, validated using dSTORM super-resolution microscopy (**Fig. 3D**).

**Figure 3.**
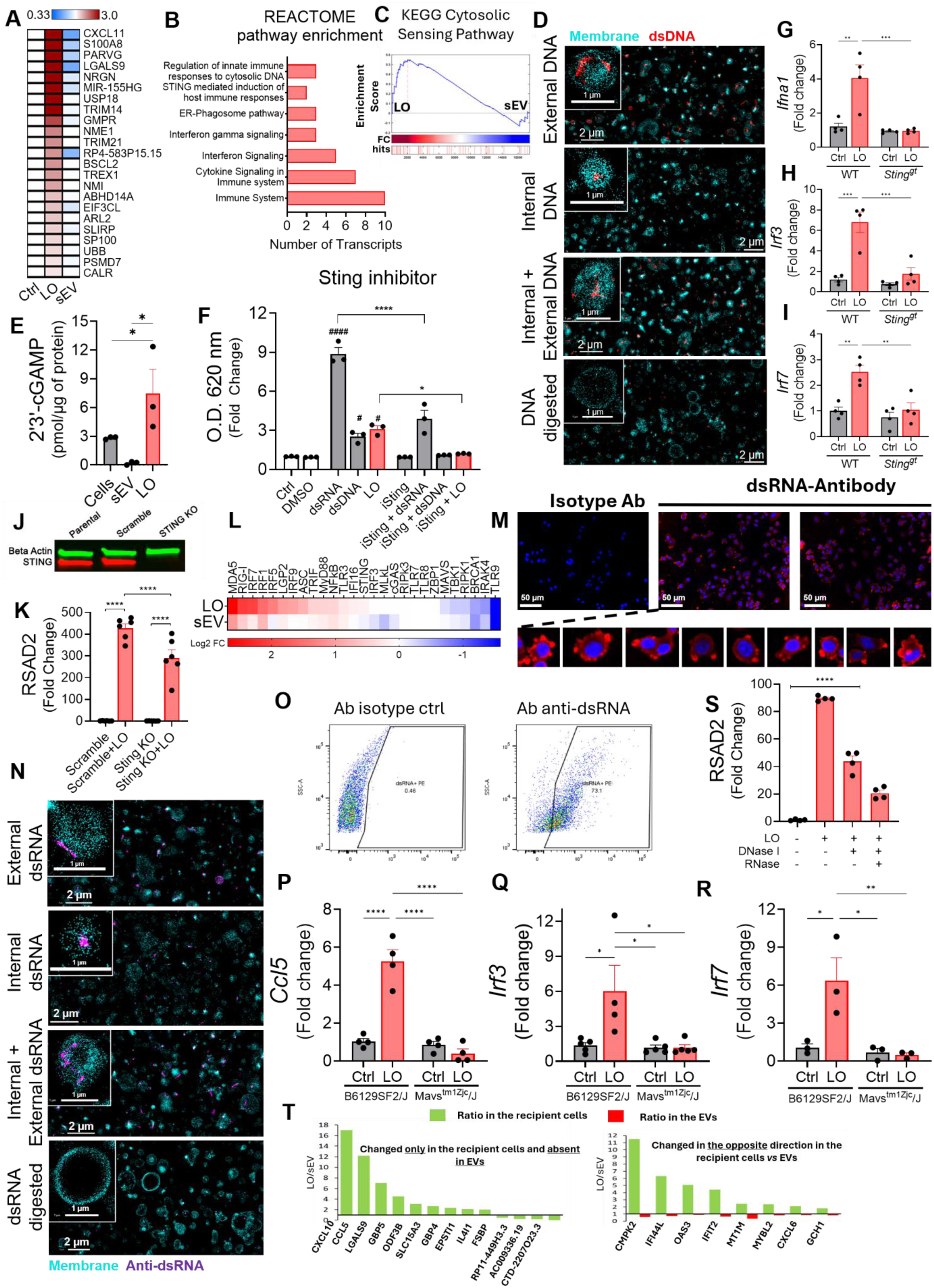
Large oncosomes engage convergent cytosolic DNA- and RNA-sensing pathways in recipient stromal cells. **A**, Heatmap of transcripts uniquely upregulated, Log2 fold change of differentially expressed, and incongruent genes between LO-treated and sEV-treated human BM-MSCs relative to vehicle (adjusted P < 0.05). **B**, Reactome pathway enrichment analysis of the gene set shown in **A**. **C**, Gene set enrichment analysis (GSEA) of the KEGG cytosolic nucleic acid–sensing pathway in LO-treated versus sEV-treated human BM-MSCs. **D**, Direct stochastic optical reconstruction microscopy (dSTORM) super-resolution imaging of PC3-derived large oncosomes labeled with a pan-EV membrane dye (cyan) and anti-dsDNA antibody (red). LO-associated DNA was assessed under four conditions: no permeabilization (External DNA), permeabilization only (internal + external DNA), DNase + permeabilization (DNA digested), and sequential DNase digestion followed by permeabilization (internal DNA). DNase treatment for Internal DNA was performed prior to permeabilization to selectively remove exposed (external) DNA while preserving membrane-protected internal DNA. Scale bars, 1 and 2 µm. **E**, Competitive ELISA quantification of 2′,3′-cGAMP in PC3 cell lysates, PC3-derived sEV, and LO. **F**, Secretable alkaline phosphatase (SEAP) reporter activity in Interferon Stimulated Genes (ISG) reporter cells treated with LO in the presence or absence of a STING inhibitor. Poly(I:C) and poly(dA:dT) were included as positive controls. **G-I**, RT–qPCR analysis of *Irf3* (**G**), *Irf7* (**H**), and *Ifna1* (**I**) expression in mouse bone marrow cells isolated from wild-type or *Sting^Gt^* mice following LO treatment for 24 h. Expression was normalized to *Actb*. **J**, Fluorescent western blot confirming STING knockout in human primary BM-MSCs generated using CRISPR–Cas9. β-actin was used as a loading control. **K**, RT–qPCR analysis of *RSAD2* expression in STING-knockout and scramble-control human BM-MSCs following LO treatment for 24 h. Expression was normalized to ACTB. **L**, Heatmap showing expression of nucleic acid–sensing pathway genes in human BM-MSCs treated with LO or sEV relative to vehicle, based on RNA-seq data. **M**, Immunofluorescence imaging of PC3 cells stained with anti-dsRNA antibody (red) or isotype control and counterstained with DAPI. Representative images and higher-magnification views are shown. **N**, dSTORM super-resolution imaging of PC3-derived LO labeled with a pan-EV membrane dye (cyan) and anti-dsRNA antibody (magenta). LO-associated dsRNA was assessed under four conditions: no permeabilization (External dsRNA), permeabilization only (internal + external dsRNA), RNase III + permeabilization (dsRNA digested), and sequential RNase III digestion followed by permeabilization (internal dsRNA). RNase III treatment for Internal dsRNA was performed prior to permeabilization to selectively remove exposed (external) dsRNA while preserving membrane-protected internal dsRNA. Scale bars, 1 and 2 µm. **O**, Flow cytometry analysis of PC3-derived LO (>1 µm) and stained with anti-dsRNA antibody or isotype control. **P-R**, RT–qPCR analysis of Ccl5 (**P**), Irf3 (**Q**), and Irf7 (**R**) expression in bone marrow cells isolated from B6;129SF2/J and B6;129-Mavs^tm1Zjc^/J (Mavs-deficient) mice and treated in vitro with PC3-derived LO. **S**, RT–qPCR analysis of *RSAD2* expression in human BM-MSCs treated with LO pre-incubated with DNase I or RNases. **T**, Comparative analysis of gene expression changes in recipient human BM-MSCs and corresponding EV preparations. Data are shown as LO/sEV ratios. Green bars indicate expression changes in recipient cells, whereas red bars indicate expression levels in EVs. **Left**, genes altered only in recipient cells and absent or negligible in EV cargo; **right**, genes showing opposite-direction changes in recipient cells versus EVs, where response to sEV < response to LO in recipient cells but expression in sEV > expression in LO in EVs. For all in vitro experiments, data represent at least three independent experiments with at least three biological and three technical replicates. Statistical analyses are described in the Methods.

We have previously reported that LOs are an EV population enriched in nucleic acid materials, when compared to sEVs. This includes diverse species of DNA and RNA^[9,15–17,24,25]^, which might be triggering the cGAS-STING activation. Moreover, surprisingly, in comparison to sEVs and even to the donor cancer cells themselves, LOs contained higher levels of 2’3’-cyclic GMP-AMP (2’,3’-cGAMP), the endogenous second messenger produced by cGAS upon DNA sensing (**Fig. 3E**). This result suggests not only active shedding of this natural STING ligand in LOs, but also that 2’,3’-cGAMP could trigger the STING signaling pathway in recipient cells even more readily than nucleic acids. Intriguingly, even if sEVs were used in a 1000-fold excess in comparison to LOs, they did not contain detectable amounts of this molecule (**Fig. 3E**), suggesting that cancer cells actively shed most of the molecule into LOs.

Using an ISG reporter system stably lentiviral transduced into BM-MSC, we found that LOs robustly activated reporter signaling at levels comparable to poly(dA:dT), a synthetic double stranded DNA used as a positive control (**Fig. 3F** and **Fig. S3A**) in recipient cells. A potent covalent small-molecule inhibitor of STING (H151) abrogated both LO- and poly(dA:dT)-induced activation of the reporter activity, corroborating the hypothesis that STING is associated with the interferon response induced by LOs (**Fig. 3F**). Interestingly, STING inhibition also significantly reduced reporter activity in cells treated with the synthetic double stranded RNA poly(I:C), suggesting a potential involvement of STING not only in dsDNA but also in dsRNA-associated signaling pathways.

To assess the contribution of STING signaling to LO-induced inflammatory activation, we first treated mixed primary mouse bone marrow cultures, which included BM-MSCs among other bone marrow cell populations, with LOs. LO treatment induced the expression of the antiviral genes Irf3, Irf7, and Ifna1 in wild-type cultures, whereas this response was markedly impaired in bone marrow cells from mice lacking functional STING (Sting1^Gt/Gt mice) (**Fig. 3G–I**). We next tested this pathway in human primary BM-MSCs by generating CRISPR-Cas9-mediated knockout of the human STING1 gene (**Fig. 3J,K**). Loss of STING attenuated, but did not completely abolish, LO-induced RSAD2 expression. These results indicate that STING signaling contributes to the inflammatory response triggered by LOs, but also suggest that additional nucleic acid-sensing pathways participate in this response, particularly in human BM-MSCs.

To further investigate this possibility, we next examined the transcriptional landscape of nucleic acid-sensing pathways induced by LOs. Further analysis of the transcriptional changes in response to PC3-derived EVs confirmed a robust induction of multiple cytosolic RNA sensors, including *MDA5, DDX58/RIG-I*, and *LGP2*, together with interferon regulatory factor family members such as *IRF1, IRF5, IRF7*, and *IRF9*, in LO-treated BM-MSCs, whereas the response to sEVs was negligible (**Fig. 3L**). GSEA further confirmed enrichment of a dsRNA-response in LO-treated cells (**Fig. S3B**).

Given these findings, we next investigated whether dsRNA was associated with LO biogenesis and cargo. PC3 cells stained with a dsRNA antibody exhibited dsRNA accumulation in correspondence of the plasma membrane blebs (**Fig. 3M**), visually confirming enrichment of this nucleic acid at sites where LOs originate. In confirmation, dSTORM imaging of LOs revealed dsRNA association with LOs both internally and externally (**Fig. 3N**). In parallel, flow cytometric characterization of LOs revealed that over 50% of particles larger than 1 µm stained positive for dsRNA, suggesting substantial accumulation of dsRNA within LOs (**Fig. 3O** and **Fig. S3C**).

To functionally evaluate the contribution of cytosolic RNA sensing to the stromal response induced by LOs, we next examined the MAVS pathway, a central downstream signaling adaptor shared by both MDA5 and RIG-I. In primary bone marrow cells obtained from B6;129SF2/J (wild-type) and B6;129-Mavs^tm1Zjc^/J (Mavs KO) mice, LO treatment induced *Ccl5, Irf3*, and *Irf7* in wild-type but not Mavs KO BM cells (**Fig. 3P-R**). Together, these results demonstrate that LO-associated dsRNA activates cytosolic RNA-sensing pathways in recipient cells and further support a model in which LOs engage redundant nucleic acid-sensing mechanisms converging on inflammatory interferon signaling.

To directly assess the contribution of LO-associated nucleic acids to this inflammatory response, we next permeabilized the LOs and enzymatically degraded DNA and RNA associated with them before treating the recipient cells. Degradation of LO-associated nucleic acids attenuated LO-induced inflammatory signaling, with DNase I treatment significantly reducing RSAD2 and CXCL8 expression in BM-MSCs and combined DNA/RNA degradation resulting in an even greater suppression (**Fig. 3S** and **Fig. S3D**).

Finally, to determine whether LO-induced transcriptional alterations were due to activation, in response to these EVs, of endogenous transcriptional programs or a mere reflection of the transfer of EV-associated mRNAs to the recipient cells, we compared RNA abundance in LOs and sEVs with the transcriptional response induced by these two EV populations in BM-MSCs. Transcripts more strongly upregulated by LOs in recipient cells were not necessarily enriched in LO themselves (**Fig. 3T**). We then compared, for each transcript, the LO/sEV expression ratio in recipient BM-MSCs with the corresponding LO/sEV ratio in the EV populations themselves. First, a substantial fraction of transcripts was altered exclusively in recipient cells while absent or negligible in EV cargo, consistent with endogenous transcriptional activation downstream of EV-mediated signaling (**Fig. 3T, left**). Second, a subset of genes displayed discordant behavior, being more strongly induced by LOs in recipient cells despite being more abundant in sEV cargo, arguing against direct RNA transfer as the primary driver of these responses (**Fig. 3T, right**). Collectively, this supports a model in which LOs primarily induce inflammatory reprogramming of BM-MSCs through activation of endogenous host transcriptional programs rather than through delivery of functional mRNA cargo.

### LO-induced IFN programming of MSCs promotes accumulation and polarization of immunosuppressive neutrophils

Having established that LOs deliver molecules that activate innate immune sensing pathways in recipient cells (**Fig. 3**), we next asked whether the response to LOs translates into functional changes in immune cell recruitment and phenotypic remodeling within the bone microenvironment. Transcriptomic analysis of LO-treated BM-MSCs revealed robust upregulation of chemotactic programs. A heatmap of the top 50 upregulated genes revealed significant enrichment of transcripts associated with leukocyte trafficking and neutrophil migration in LO-treated cells, a pattern that was largely absent in sEV treated cells (**Fig. 4A**). Consistently, GSEA demonstrated strong enrichment of the GO neutrophil chemotaxis signature in LO-treated BM-MSCs compared to sEVs (NES = 2.66) and control (NES = 3.71) (**Fig. 4B-C**)

**Figure 4.**
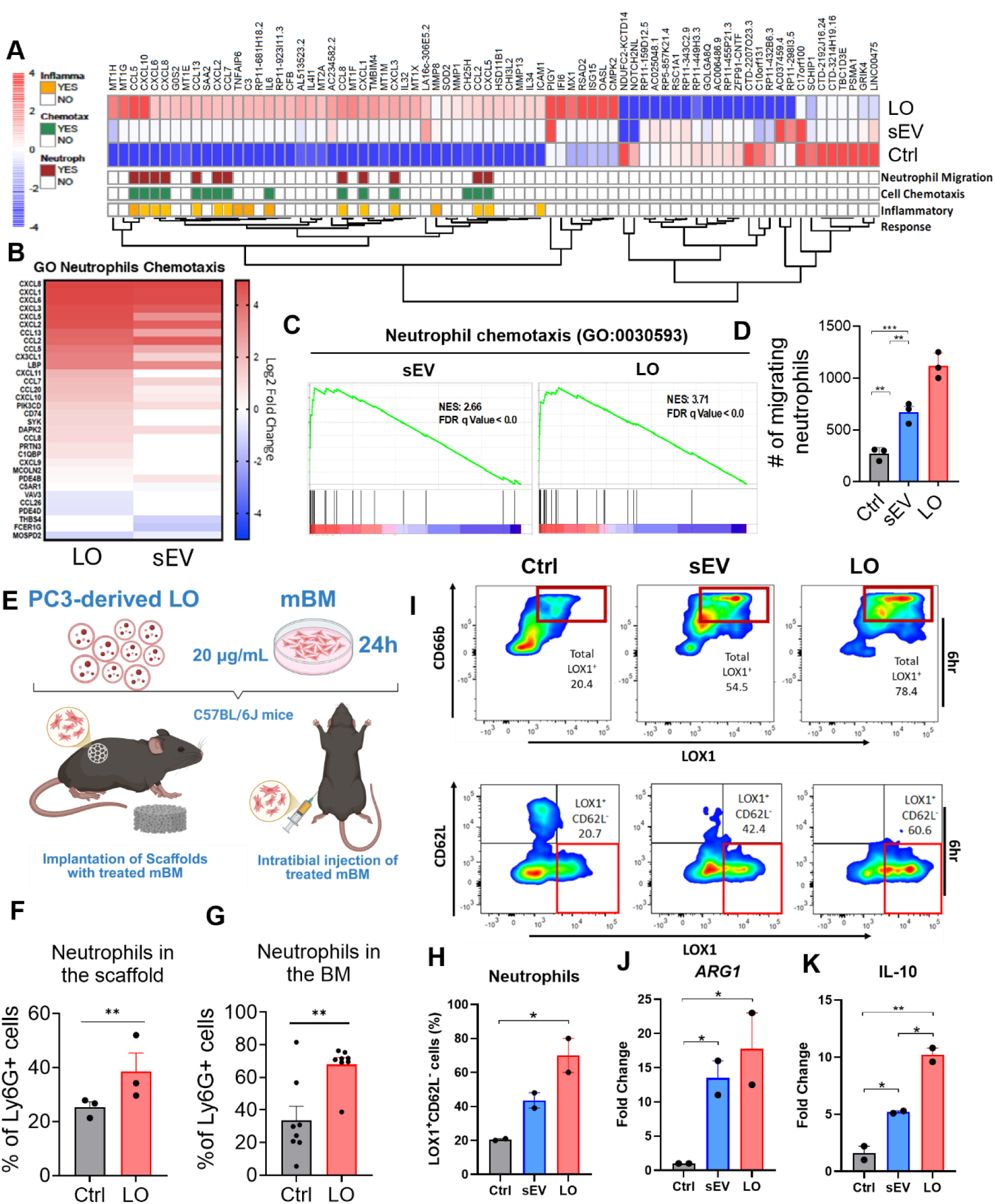
LO-induced reprogramming of BM-MSCs promote recruitment and differentiation of neutrophils into PMN-MDSC phenotype. **A**, Heatmap of the top 50 upregulated genes in human bone marrow–derived mesenchymal stem cells (BM-MSCs) treated with large oncosomes (LO) relative to vehicle control, based on the RNA-seq dataset described in Fig. 2A (n = 3 biological replicates per group). **B-C**, Gene set enrichment analysis (GSEA) of the Gene Ontology neutrophil chemotaxis pathway (GO:0030593) in LO-treated or sEV-treated versus vehicle-treated human BM-MSCs, ranked by log2 fold change. **D**, Transwell migration assay quantifying recruitment of primary human neutrophils freshly isolated from peripheral blood toward conditioned medium from vehicle-, sEV- or LO-treated human BM-MSCs. Each dot represents one independent experiment (n=3/group). **E**, Experimental schematic of neutrophil infiltration into either subcutaneous scaffolds seeded with vehicle- or LO-treated mouse bone marrow (mBM) and implanted subcutaneous or vehicle- or LO-treated mBM injected intratibially (n=4/group) into C57BL/6J mice. Scaffolds were harvested after 1 week (n = 3 mice per group) post implantation and bone marrows harvested 48h after injection. **F-H,** quantification of the percentage of Ly6G^+^ from the total CD45^+^ cells in the scaffold (**F**) and bone marrow (**G**). **H-I**, Flow cytometry analysis of primary human neutrophils exposed for 6 h to conditioned medium from vehicle-, sEV- or LO-treated human BM-MSCs, assessing CD62L, CD66b and LOX-1 expression. **J-K**, RT–qPCR analysis of ARG1 (**J**) and IL10 (**K**) expression in primary human neutrophils following 6 h exposure to conditioned medium from vehicle-, sEV- or LO-treated BM-MSCs. Expression was normalized to ACTB. Each dot represents one independent experiment.

To assess the functional impact of this transcriptional remodeling, we performed transwell migration assays using conditioned medium from EV-treated BM-MSCs, confirming a significantly increased neutrophil migration on LO-treated compared to the vehicle- or sEV-treated cells conditioned medium (**Fig. 4D**). To validate these observations *in vivo*, we employed two complementary approaches. LO-educated C57BL/6J bone marrow cells were either embedded within 3D polyurethane scaffolds^[35]^ and implanted subcutaneously into syngeneic mice (**Fig. 4E,F**) or injected directly into the tibiae of syngeneic recipients (**Fig. 4E,G**). In both cases, we observed a significant increase of Ly6G⁺ neutrophils in response to the LO-treated cells.

We next examined whether LOs might modulate the neutrophil phenotype. Human neutrophils exposed to conditioned medium from LO-treated BM-MSCs displayed a striking increase in Lectin-type oxidized LDL receptor 1 (LOX-1) and CD66b expression and near-complete loss of CD62L. This result was time-dependent and visible even after just one hour if exposure to the conditioned medium (**Fig. 4H-I** and **S4**). LOX-1+ neutrophils are a specialized immature subset of neutrophils that function as Polymorphonuclear Myeloid-Derived Suppressor Cells (PMN-MDSCs) that are known to heavily suppress T-cell immune responses in cancer^[41]^. Upregulation of the immunosuppressive genes ARG1 and IL10 (**Fig. 4J-K**) further suggested a functional polarization toward a PMN-MDSC-like state.

### Tumor-derived LOs are necessary and sufficient for systemic accumulation of PMN-MDSCs and tumor progression

Building on our findings that LO-induced programming of BM-MSCs promotes neutrophil accumulation and PMN-MDSC-like polarization (**Fig. 4**), we next asked whether tumor-derived LOs are required to drive this systemic immunosuppressive neutrophil response *in vivo*, by developing an orthotopic breast cancer model with Septin 2 (Sept2) knockout to impair LO shedding (**Fig. 5A**). Sept2 is a conserved GTP-binding cytoskeletal protein implicated in membrane blebbing and cortical organization in cancer cells, including bleb-associated signaling in detached melanoma cells^[42]^, although its involvement in LO biogenesis has not been previously tested. Efficient loss of Sept2 following CRISPR-Cas9-mediated editing was confirmed by immunoblotting across independent monoclonal 4T1.2 populations (**Fig. 5B**). Notably, we observed a marked reduction in the release of large particles from Sept2-deficient cells compared with scramble controls (**Fig. 5C**), identifying Sept2 as a potential regulator of LO shedding. BALB/c mice were then implanted with either WT or Sept2 KO 4T1.2 cells, while additional matched cohorts received systemic LO injections as a rescue strategy (**Fig. 5A**). Tumor engraftment and growth were comparable between groups, as assessed by caliper measurements (**Fig. S5A**). dSTORM analysis of plasma-derived EVs revealed a significant reduction in circulating PIGS⁺ and Caveolin-1⁺ large particles in mice bearing Sept2 KO tumors, confirming the ability of Sept2 loss to suppress LO abundance *in vivo*.

**Figure 5.**
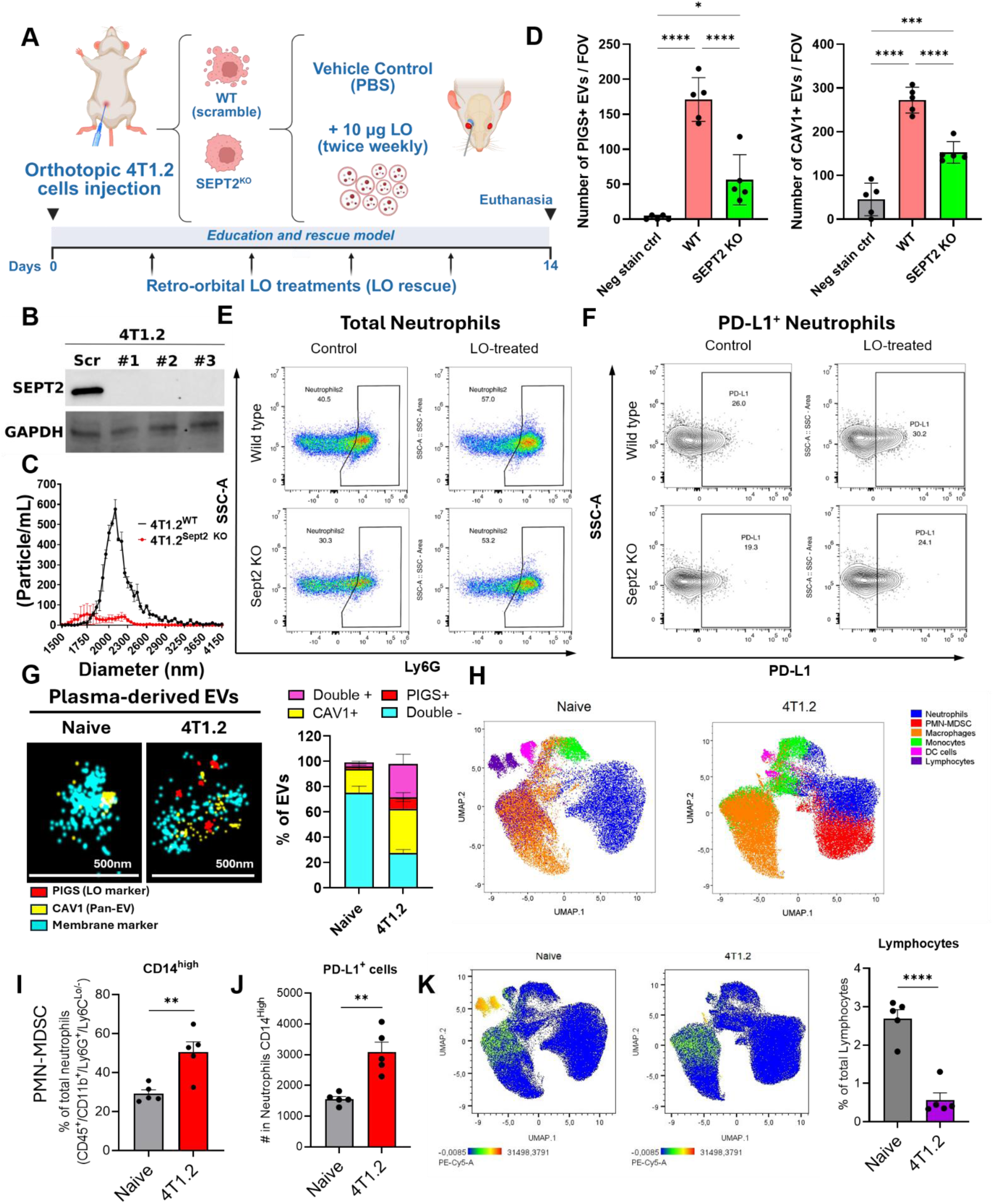
LO-induced reprogramming of BM-MSCs induce accumulation and polarization of neutrophils into PMN-MDSC phenotype. **A**, schematic representation of the LO-deficient and LO-rescue orthotopic tumor model developed. Balb/c mice were orthotopic injected into the mammary fat pad with 4T1.2 cells engineered to lack Sept2 (LO deficient) or not. A separate experimental group set was performed including systemic injection of LO to rescue LO-induced signaling in absence of sept2. **B,** SEPT2 western blot analysis of scramble control and 3 individual monoclonal populations of 4T1.2 Sept2 KO induced using CrisprCas9. **C,** microfluidic-resistive pulse sensing particle count analysis quantifying LO shedding in scramble control and Sept2 KO 4T1.2 cell lines. **D,** quantification of the total number of particles per field of view (FOV) positive for PIGS (LO marker) and Caveolin 1 (Pan-EV marker) in the plasma from the tumor-bearing mice described in Figure 5A. **E-F**, spectral flow cytometry analysis of the bone marrow of the mice in Figure 5A, showing the percentage of Ly6G+ cells (neutrophils) (**E**) and the percentage of PD-L1+ neutrophils (PMN-MDSC) (**F**). **G**, dSTORM analyses of particles isolated from the plasma of BALB/c mice naïve vs orthotopically implanted with the bone metastatic 4T1.2 breast cancer cell line with high LO shedding. Imaging (left) and quantification (right) of the percentage of particles positive or negative for either PIGS (red) or Caveolin 1 (yellow). A general membrane dye was used for counterstaining (cyan). **H**, UMAP visualization and FlowSOM clustering of high-parameter spectral flow cytometry data from bone marrow cells of naïve and 4T1.2 tumor-bearing mice. Data represents concatenated files from all animals (n = 5 mice per group). **I**, quantification of PMN-MDSC in the flow cytometry analysis from panel **6H**. **J**, PD-L1 expression within the PMN-MDSC cluster identified in panel **6I**. **K**, UMAP visualization (left) and quantification (right) of CD3 expression across bone marrow immune populations from naïve and tumor-bearing mice. Statistical analyses are described in the Methods.

We next investigated how tumor-derived LO production shapes the composition of bone marrow myeloid populations. Neutrophils and PMN-MDSCs were identified using a sequential gating strategy based on CD45, CD11b, Ly6G, and PD-L1 expression (**Fig. S5C**). Spectral flow cytometry revealed a reduction in the overall frequency of Ly6G⁺ neutrophils in mice bearing Sept*2* KO tumors relative to controls (**Fig. 5E**). More strikingly, the proportion of PD-L1⁺ neutrophils, consistent with a PMN-MDSC phenotype, was significantly decreased in the absence of tumor-derived LOs (**Fig. 5F**). Importantly, in a LO-rescue model, systemic administration of purified syngeneic LOs restored PD-L1^+^ neutrophil accumulation within the BM compartment, suggesting that LOs are indeed sufficient to drive this immunosuppressive phenotype *in vivo* (**Fig. 5F**).

To investigate how prolonged tumor progression influences the BM compartment, we compared healthy and tumor-bearing mice using the highly metastatic, LO-producing 4T1.2 orthotopic breast cancer model. BALB/c mice were monitored for four weeks following mammary fat pad implantation, after which bone marrow was collected for flow cytometric analysis (**Fig. S5B**). Super resolution dSTORM imaging confirmed a substantial increase in the proportion of circulating PIGS⁺ large particles in mice bearing orthotopic 4T1.2 tumors compared to naïve controls (**Fig. 5G**). High-dimensional spectral flow cytometry, combined with UMAP projection and FlowSOM clustering, revealed a pronounced expansion of neutrophil populations within the BM of tumor-bearing animals (**Fig. 5H**). More specifically, we observed a significant increase in the frequency of cells with high expression of CD14, CXCR2, and PD-L1, and reduced expression of MHC-II (**Fig. 5I-J** and **S5C-E**), compatible with a PMN-MDSC phenotype.

Notably, PMN-MDSC abundance in the BM positively correlated with primary tumor size, indicating a link between tumor growth and BM expansion of this immunosuppressive population (**Fig. S5F**). BM cells further exhibited a marked reduction in CD3⁺ lymphoid populations (**Fig. 5K**), consistent with the establishment of an immunosuppressive niche. These observations occurred in the absence of detectable GFP⁺ tumor cells in the BM, suggesting that the LO-induced immunological reprogramming precedes metastatic colonization. Notably, no significant changes were observed in macrophage populations (**Fig. S5G-H**), indicating that the LO-driven remodeling identified in this study, which results in the upregulation of cytokines known to regulate myeloid cells, preferentially impacts the neutrophil compartment *in vivo*.

Together, these data identify a previously unrecognized mechanism by which tumor-derived LOs shape the pre-metastatic niche. By reprogramming bone marrow stromal cells toward a pro-inflammatory, neutrophil-accumulating state, LOs drive the polarization of immunosuppressive PMNs while promoting lymphocyte depletion. These findings, collectively, establish LOs as potent systemic mediators of metastatic niche formation and provide a mechanistic link between tumor-derived vesiculation, stromal reprogramming, and early immune remodeling at distant sites.

## DISCUSSION

LOs have emerged as a distinctive class of tumor-derived EVs associated with metastatic and lethal cancer^[11,12]^, yet their functional role in shaping metastatic progression has remained incompletely defined. In this study, we provide evidence that LOs act as potent systemic mediators of pre-metastatic niche formation by coupling tumor-intrinsic vesiculation programs to innate immune activation in stromal cells. By integrating cargo characterization, transcriptomic profiling, and *in vivo* functional models, we establish LOs not merely as enlarged EVs, but as specialized and highly complex signaling platforms that orchestrate inflammatory and immunosuppressive remodeling of the bone microenvironment.

Mechanistically, this study identifies cytosolic nucleic acid sensing as the central pathway underlying LO-induced stromal activation. LOs have emerged as the EV population enriched in nucleic acids compared with sEVs^[12,16,17,21,25]^. In our findings, this translates into an unexplored mechanism of intercellular communication where distinct nucleic acid species converge on redundant signaling pathways, specifically STING for DNA and MAVS (MDA5/RIG-I) for dsRNA, within recipient cells. The convergence of DNA and RNA sensing pathways suggests that LOs act as composite carriers of immunostimulatory signals capable of activating a coordinated innate immune response. Consistent with this concept, recent work has shown that epigenetic reprogramming through EZH2 inhibition induces accumulation of endogenous dsRNA and activates a dsRNA-STING-interferon signaling axis in prostate cancer models^[43]^.

In addition, a particularly intriguing observation is the high enrichment of 2’3’-cGAMP within LOs. As a direct activator of STING, cGAMP allows LOs to propagate innate immune signaling constitutively and independently of upstream DNA sensing in recipient cells, thereby amplifying and accelerating immune activation in distant compartments. This suggests a potential mechanistic basis by which LO-derived nucleic acids may elicit stronger interferon responses than those delivered by sEVs. Such a mechanism may be especially relevant in the context of metastatic dissemination, where rapid and robust reprogramming of the microenvironment is required to support tumor cell colonization^[44]^.

Findings from this study also define a stromal-mediated axis through which tumor-derived vesicles indirectly shape immune composition, linking vesicle signaling to the expansion of immunosuppressive myeloid populations. PMN-MDSCs are increasingly recognized as central mediators of metastatic immune remodeling, but the upstream tumor-derived signals that promote their accumulation in distant organs remain incompletely defined. Our findings support a model in which LOs act as systemic messengers that reprogram the bone marrow microenvironment before metastatic expansion. Across complementary *in vitro* and *in vivo* approaches, LO exposure was associated with activation of a pro-inflammatory stromal program, increased production of neutrophil- and myeloid-recruiting mediators, accumulation and polarization of immunosuppressive PMN-MDSC populations, and concomitant depletion of lymphoid cells in the bone marrow. These observations suggest that LOs do not merely promote tumor cell-intrinsic metastatic traits but also remodel distant stromal and immune compartments to create a permissive pre-metastatic niche.

Interestingly, a recent study demonstrated that mitochondrial DNA released by senescent tumor cells enhances PMN-MDSC-driven immunosuppression through activation of the cGAS-STING pathway^[28]^. This mechanism may plausibly be mediated by LOs, as we have previously reported using a multiomics profiling that LOs are enriched in mitochondrial content (e.g., MT-CO1, TOMM40)^[9]^, suggesting a potential route for the transfer of immunostimulatory mitochondrial DNA to recipient cells. These discoveries extend prior models of “seed and soil” crosstalk. While previous work linked LOs to local prostate fibroblast reprogramming via AKT1-MYC^[20]^, this work uncovers a long-range axis targeting the stroma at the most frequent metastatic site for both prostate and breast cancer.

Our findings also provide important context for the role of IFN signaling in cancer. While IFN pathways are classically associated with anti-tumor immunity, accumulating evidence suggests that chronic or spatially restricted activation can promote immunosuppressive and pro-tumorigenic effects^[45,46]^. The sustained IFN response induced by LOs in stromal cells appears to drive chemokine production and myeloid cell recruitment in a manner that ultimately favors tumor progression. This highlights the context-dependent nature of innate immune signaling and suggests that tumor-derived vesicles may exploit these pathways to reprogram the host environment.

From a translational perspective, our study identifies multiple potential points of therapeutic intervention that could be further investigated in follow-up studies. Targeting LO biogenesis, release, or uptake could be a potential mechanism to disrupt early steps in metastatic niche formation. In addition, the unique molecular composition of LOs, including enriched surface markers and nucleic acid cargo^[9,16]^, may provide opportunities for biomarker development or selective targeting strategies. Intercepting circulating LOs could represent a novel strategy to disrupt tumor-host communication before metastatic colonization occurs. By limiting LO-driven conditioning of distant tissues, such approaches may help prevent pre-metastatic niche formation and potentially improve the effectiveness of subsequent systemic therapies, including immunotherapy. Also, given the systemic nature of LO-mediated signaling, such approaches may be accessed in complement with existing therapies aimed at the primary tumor or established metastases.

Although our data demonstrate a central role for LO-associated nucleic acids in driving stromal activation, the relative contributions of DNA, RNA, and cGAMP require further investigation. In addition, while we observe consistent biological responses across multiple experimental systems, including human and murine models, validation in patient-derived samples will be essential to establish clinical relevance. Finally, the mechanisms governing LO uptake, intracellular trafficking, and cargo release in recipient cells remain to be fully elucidated and represent important areas for future investigation.

In summary, we propose a model in which tumor-derived LOs function as systemic signaling hubs that deliver immunostimulatory nucleic acids and second messengers to stromal cells, activating innate immune pathways and driving the formation of an inflammatory environment which, through the acquisition of an immunosuppressive polarization counterbalance, can culminate in the formation of a pre-metastatic niche.

Through this mechanism, LOs link tumor cell plasticity and vesiculation to long-range immune remodeling, thereby facilitating metastatic progression. These findings redefine the functional landscape of tumor-derived EVs and position LOs as potential mediators of tumor-host communication in cancer metastasis.

## ACKNOWLEDGMENTS

We thank Karen Cavassani, Neil Bhowmick, David Lyden and Leigh Ellis for their valuable scientific discussions and thoughtful feedback throughout this work. We are also grateful to Ágnes Kittel for the support with the transmission electron microscopy analysis. We acknowledge the Cedars-Sinai Health Sciences University Flow Cytometry Shared Resource Core for technical support with flow cytometry and spectral flow cytometry experiments. This work was supported by the National Institutes of Health (R01CA218526 and R01CA234557 to D.D.V. and R21CA299812 to T.F.S). and by the Department of Defense Prostate Cancer Research Program (HT9425-25-1-0734 to T.F.S.).

## DATA AVAILABILITY STATEMENT

The raw dataset generated for the RNA sequencing analysis has been deposited in the NCBI Gene Expression Omnibus (GEO) under accession number GSE341387. Processed count table dataset is available as **Supplementary table 1**. All other data supporting the findings of this study are available within the article and its Supplementary Information files. Additional data and materials are available from the corresponding authors upon reasonable request.

## CONFLICT OF INTEREST

The authors declare that they have no competing interests.

## SUPPLEMENTARY FIGURES

**Supplementary Figure 1.**
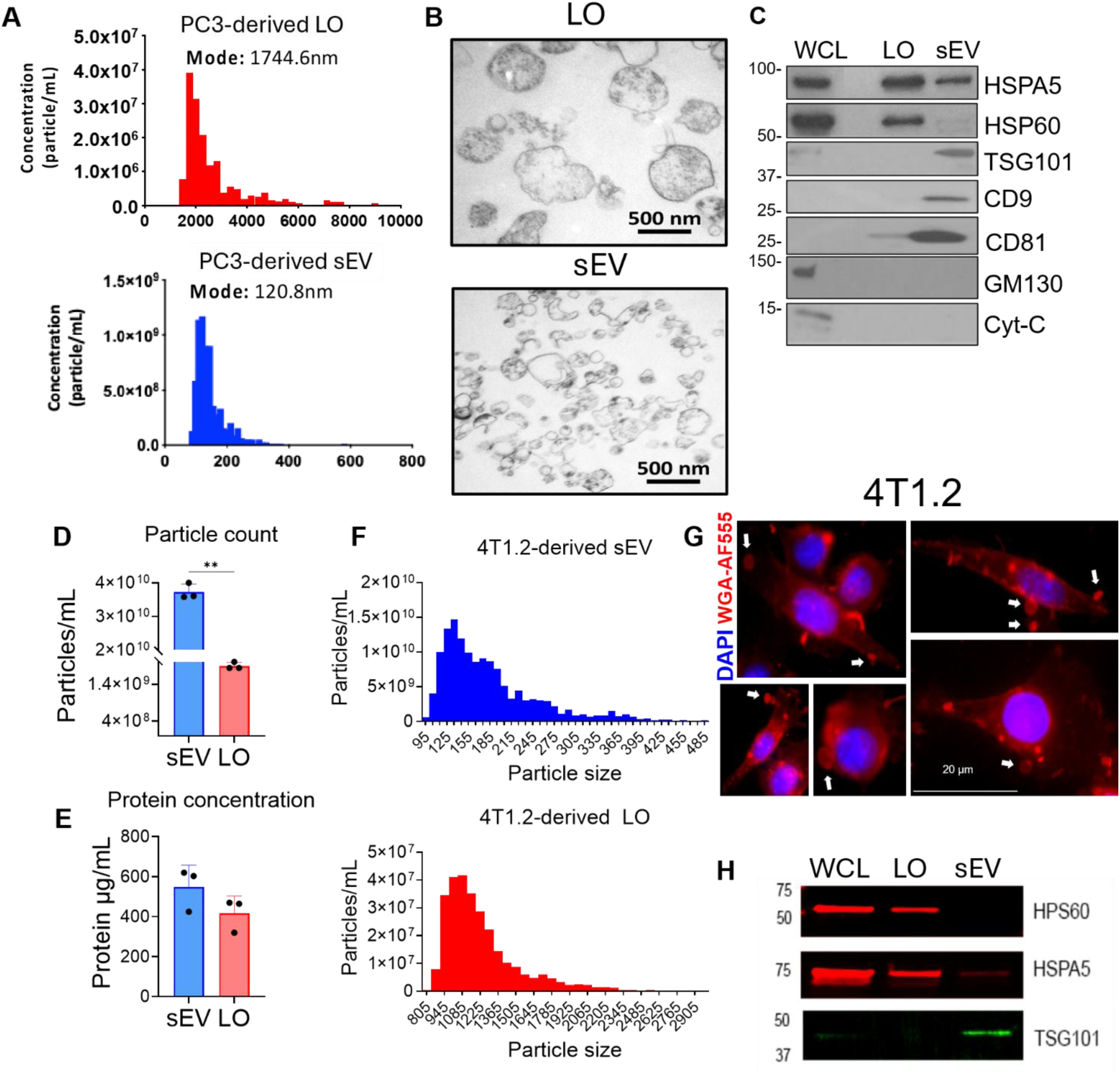
Characterization of large oncosomes and small EVs across cell lines. **A**, Tunable-resistive pulse sensing (TRPS) size-distribution histograms of PC3-derived LO (top) and sEV (bottom). **B**, Transmission electron microscopy of PC3-derived LO and sEV preparations. **C**, Western blot analysis of PC3 EVs showing LO markers (HSP60, HSPA5), sEV markers (TSG101, CD9, CD81) and cell-associated markers (GM130 and cytochrome c) using HRP-conjugated secondary antibodies. **D**, TRPS particle concentration measurements of 4T1.2-derived LO and sEV. **E**, Total protein abundance of 4T1.2-derived LO and sEV preparations. **F**, TRPS size-distribution histograms of 4T1.2-derived LO (top) and sEV (bottom). **G**, Fluorescence imaging of 4T1.2 bone-metastatic breast cancer cells showing LO shedding. Plasma membrane labeled with WGA-AF555 (red) and nuclei with DAPI (blue). Scale bar, 20 µm. **H**, Fluorescent Western blot of 4T1.2-derived EVs and whole-cell lysate showing LO markers (HSP60, HSPA5) and the sEV marker TSG101 using secondary antibodies conjugated with StarBright Blue 520 in green (anti-mouse) or StarBright Blue 700 in red (anti-rabbit).

**Supplementary Figure 2.**
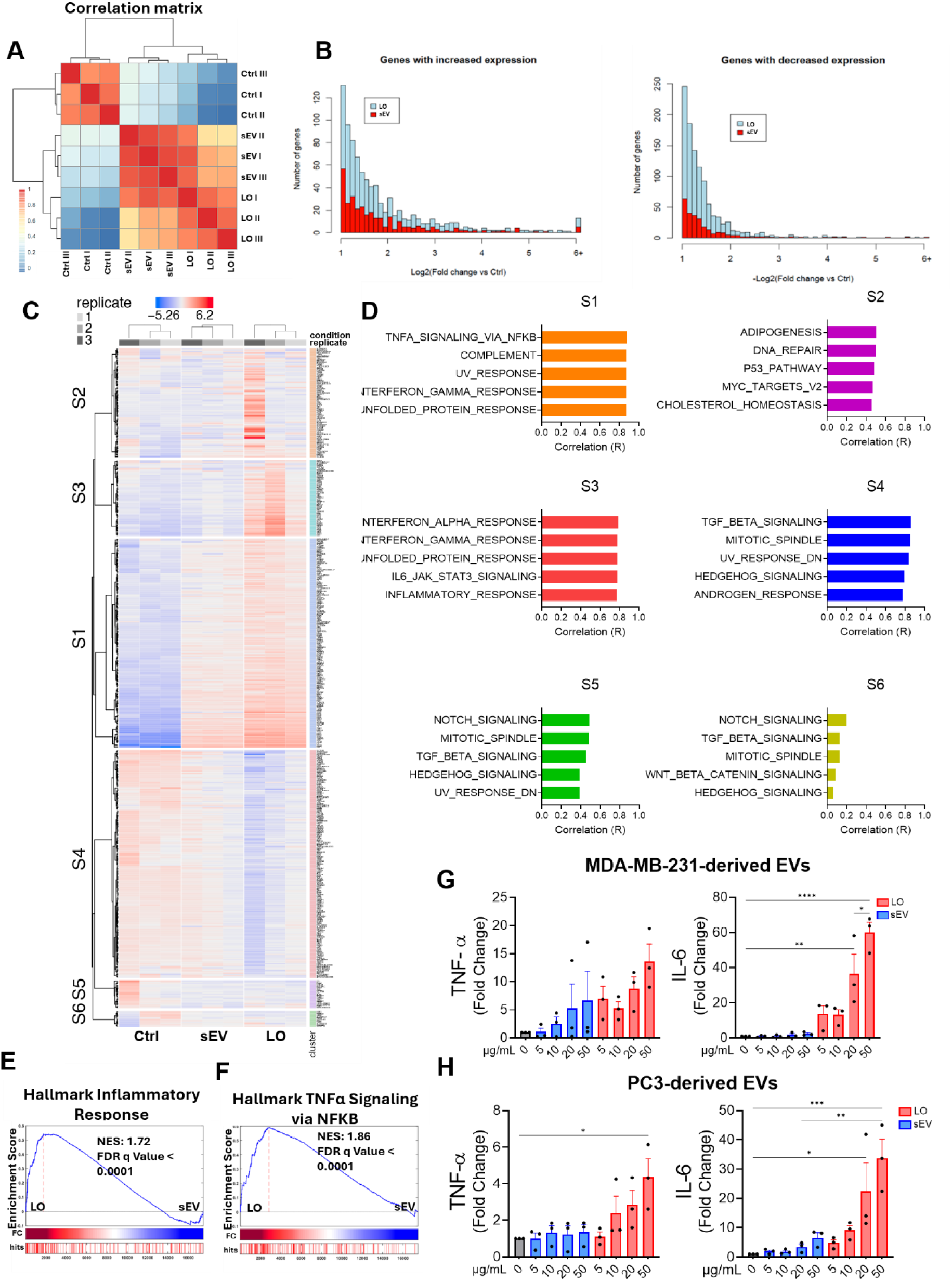
RNA-seq quality control and pathway analysis of EV-treated human BM-MSCs. **A-B**, RNA-seq quality control analyses, including sample-to-sample Pearson correlation matrix (**A**) and distributions of gene expression changes and concordance/discordance of fold-change directionality between LO and sEV treatments (**B**). **C**, Clustered Heatmap is a 2-way unsupervised hierarchical clustering technique that simultaneously clusters the expression matrix along rows and columns, clustering similar genes and similar samples together. The tree-like dendrogram shows the ‘distance’ between features and the approximate groups. The column annotations show the correlation with the phenotypes. **D**, top 5 HALLMARK gene set pathways enriched in each cluster identified in panel C. **E-F**, GSEA enrichment plots for inflammatory response (**E**) and TNFα signaling via NF-κB (**F**) in LO-treated versus sEV-treated human BM-MSCs. **G-H**, RT–qPCR analysis of IL6 and TNFA expression in human BM-MSCs treated with increasing concentrations of MDA-MB-231-derived (**G**) or PC3-derived (**H**) LO or sEV. Expression was normalized to ACTB. Each dot represents one independent experiment. Each dot represents one independent experiment. For all in vitro experiments, data represents at least three independent experiments with at least three biological and three technical replicates. Statistical analyses are described in the Methods.

**Supplementary Figure 3.**
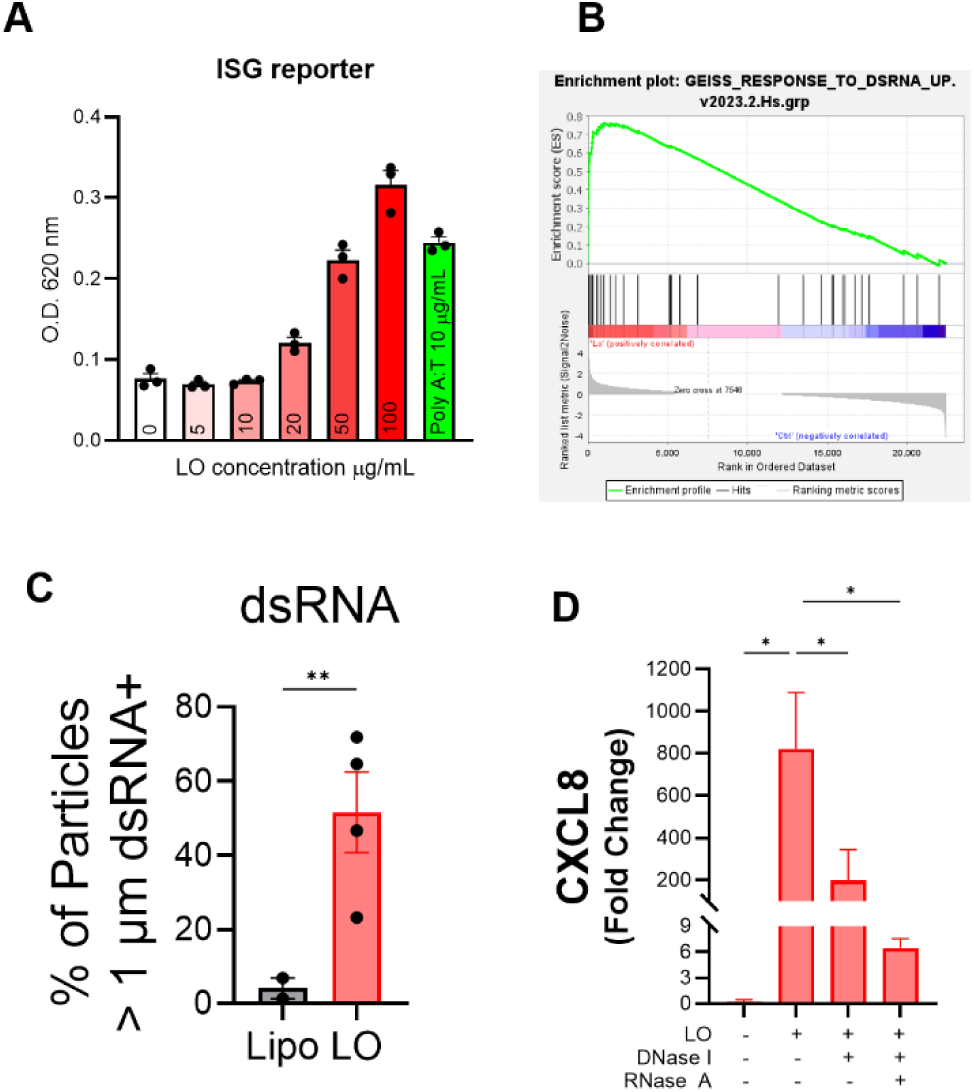
Validation and extended analyses of LO-mediated nucleic acid sensing. **A**, BM-MSC cells expressing secretable alkaline phosphatase (SEAP) under the minimal promoter of Interferon Stimulated Genes (ISG) as reporter system. Treatment with increasing LO concentrations showing correlative effect on the activity of the reporter. Poly(dA:dT) was included as a positive control. **B**, Single-cell RNA sequencing of bone marrow cells identifying BM-MSC populations among the isolated cells used in Fig. 3G**-I****. C**, Flow cytometry gating strategy and additional representative analysis of dsRNA-positive LO corresponding to Fig. 3N**. D**, GSEA of the GEISS response-to-dsRNA gene set in LO-treated versus vehicle-treated human BM-MSCs from the experiment described in Fig. 2A**. E**, RT–qPCR analysis of CXCL8 expression in human BM-MSCs treated with LO pre-incubated with DNase I or RNases, corresponding to Fig. 3S. Expression was normalized to *ACTB*.

**Supplementary Figure 4.**
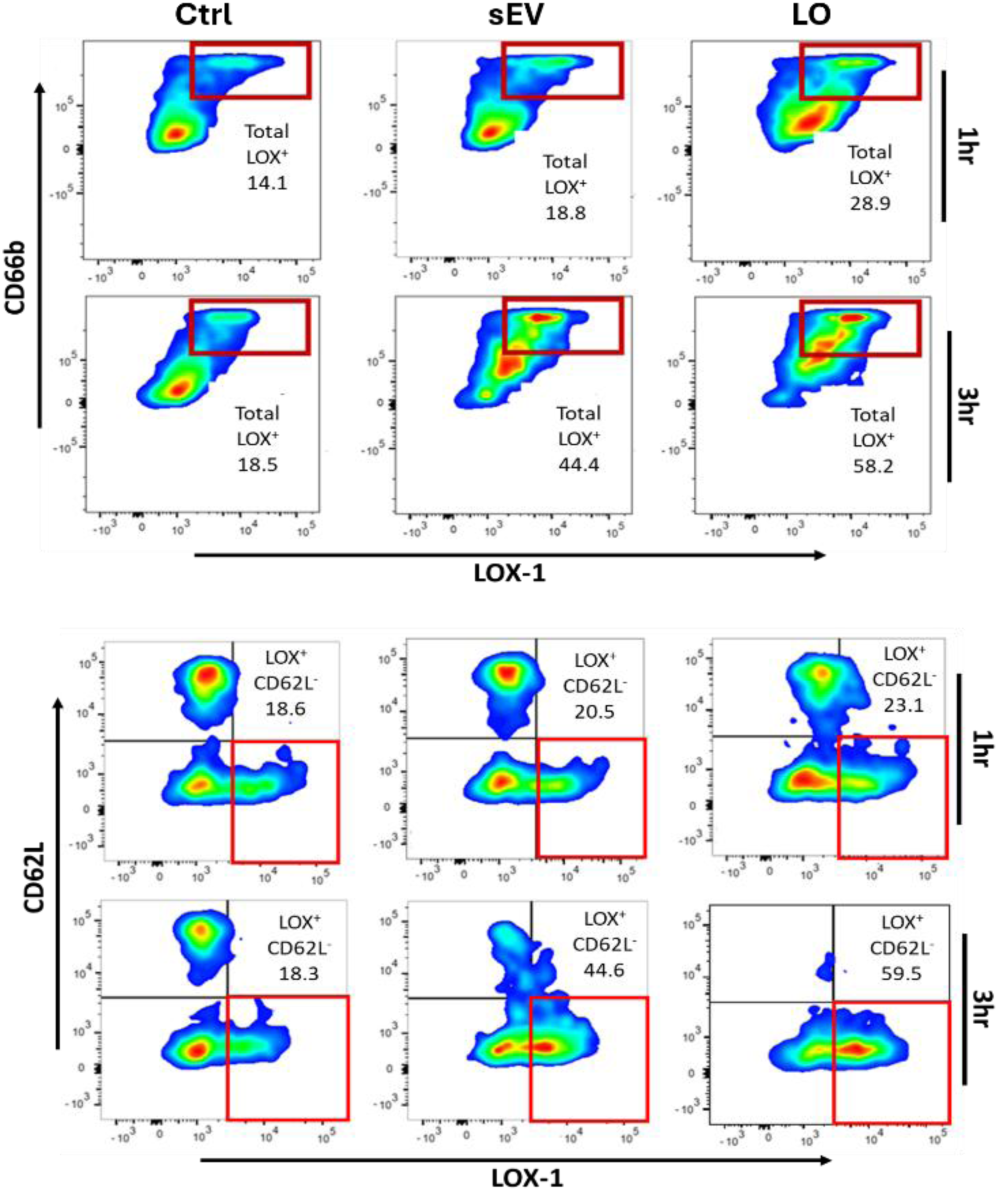
LO-treated BM-MSCs induce fast neutrophil polarization. Flow cytometry analysis of CD62L, CD66b, and LOX-1 expression in primary human neutrophils following exposure to conditioned medium from vehicle-, sEV- or LO-treated BM-MSCs at 1 and 3 h, associated with Figure 4H-I.

**Supplementary Figure 5.**
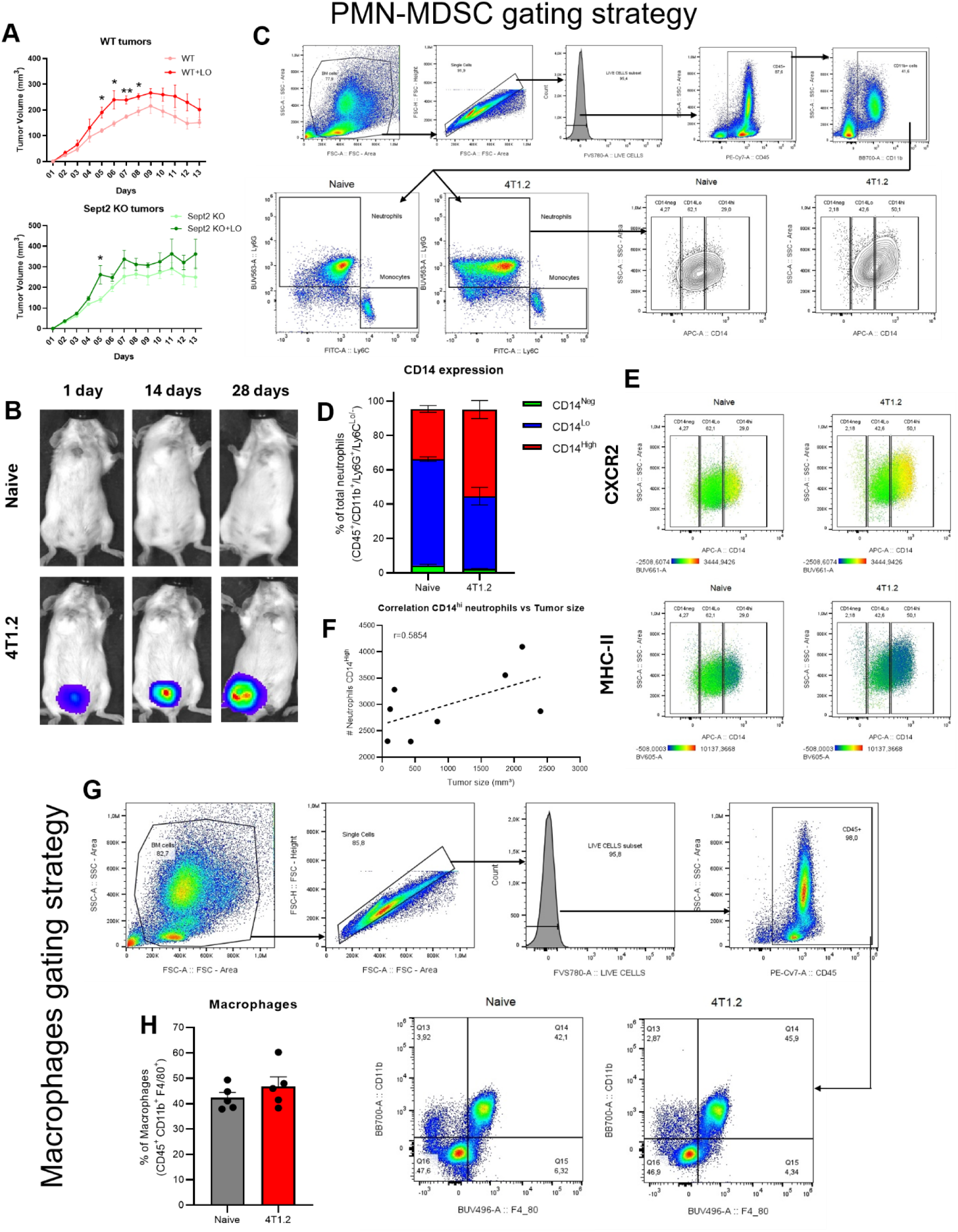
Tumor tracking and gating strategies. **A**, tumor growth measurements from experiment described in Figure 5A. **B**, Representative IVIS images of tumor growth in BALB/c mice orthotopically implanted with 4T1.2-Luc/GFP cells at days 1, 14 and 28 after implantation. Associated with Figures 5G-K and **S5C-G. C**, Gating strategy for identification of neutrophils and PMN-MDSCs based on CD45, CD11b, Ly6C, Ly6G and CD14 expression. **D**, Quantification of CD14^-^, CD14^lo^ and CD14^hi^ neutrophils in bone marrow from naïve and tumor-bearing BALB/c mice (n = 5 mice per group). **E**, Expression of CXCR2 (top) and MHC class II (bottom) across neutrophil populations with different CD14 expression level identified by spectral flow cytometry. **F**, Correlation between tumor size and frequency of CD14^hi^ neutrophils in tumor-bearing mice. **G**, Gating strategy and quantification of macrophage populations in bone marrow from naïve and tumor-bearing mice. **H**, Quantification of macrophage populations in **G**.

## TABLE LEGENDS

**Supplementary Table 1. Gene expression count matrix from RNA-seq of BM-MSCs treated with large oncosomes or small extracellular vesicles.** Gene-level RNA-seq counts from bone marrow mesenchymal stem cells (BM-MSCs) treated with PC3-derived large oncosomes (LOs) or small extracellular vesicles (sEVs) for 24 h. Rows represent individual genes and columns represent biological samples used for downstream differential gene expression analyses.

## Notes

### Competing Interest Statement

The authors have declared no competing interest.

